# Systematic assessment of commercially available low-input miRNA library preparation kits

**DOI:** 10.1101/702456

**Authors:** Fatima Heinicke, Xiangfu Zhong, Manuela Zucknick, Johannes Breidenbach, Arvind Sundaram, Siri T. Flåm, Magnus Leithaug, Marianne Dalland, Andrew Farmer, Jordana M. Henderson, Melanie A. Hussong, Pamela Moll, Loan Nguyen, Amanda McNulty, Jonathan M. Shaffer, Sabrina Shore, HoiChong Karen Yip, Jana Vitkovska, Simon Rayner, Benedicte A Lie, Gregor D. Gilfillan

## Abstract

High-throughput sequencing is increasingly favoured to assay the presence and abundance of micro RNAs (miRNAs) in biological samples, even from low RNA amounts, and a number of commercial vendors now offer kits that allow miRNA sequencing from sub-nanogram (ng) inputs. However, although biases introduced during library preparation have been documented, the relative performance of current reagent kits has not been investigated in detail. Here, six commercial kits capable of handling <100ng total RNA input were used for library preparation, performed by kit manufactures, on synthetic miRNAs of known quantities and human biological total RNA samples. We compared the performance of miRNA detection sensitivity, reliability, titration response and the ability to detect differentially expressed miRNAs. In addition, we assessed the use of unique molecular identifiers sequence (UMI) tags in one kit. We observed differences in detection sensitivity and ability to identify differentially expressed miRNAs between the kits, but none were able to detect the full repertoire of expected miRNAs. The reliability within the replicates of all kits was good, while larger differences were observed between the kits, although none could accurately quantify the majority of miRNAs. UMI tags, at least within the input ranges tested, offered little advantage to improve data utility. In conclusion, biases in miRNA abundance are heavily influenced by the kit used for library preparation, suggesting that comparisons of datasets prepared by different procedures should be made with caution. This article is intended to assist researchers select the most appropriate kit for their experimental conditions.

## Introduction

Micro RNAs (miRNAs) are ~22 nucleotide long non-coding small RNAs that regulate gene expression at a post-transcriptional level by binding to their mRNA targets to inhibit translation. First discovered in the early 1990s^1 2^, miRNAs have been shown to impact biological processes such as cellular differentiation and development^3–6 7–9 10–12 13 14^ Alterations in miRNA expression have been observed in various diseases^15 16 17^ and an accurate method for detecting and measuring miRNA expression is therefore crucial. In recent years, next generation sequencing (NGS) has evolved as the method of choice. The main advantages of NGS, compared to qPCR and microarray techniques, are the possibility to discover novel miRNAs and the ability to detect differences in miRNA sequences on a single base level. Furthermore, NGS enables the study of low-abundance miRNAs, which is especially useful when examining miRNAs in specific cell types or body fluids like serum and plasma. Accordingly, the latest miRNA library preparation kits allow inputs as low as 100 picograms total RNA. The library preparation process typically consists of (i) addition of adapter sequences onto the miRNA, (ii) reverse transcription and (iii) PCR amplification prior to sequencing. The kits investigated in this study used both two adapter and singleadapter circularization protocols which can broadly be divided into two classes: those employing RNA ligases (e.g. T4 RNA ligase) and those employing polyadenylation (poly-A) and template-switching oligonucleotides to attach adapter sequences to the single-stranded miRNAs.

Despite the reported advantages of NGS, the miRNA abundance detected by sequencing and that in the original sample have been shown to differ by up to four orders of magnitude^18^. In particular, the addition of adapters onto the miRNA insert has been identified as a major contributor to this bias^19 20^. For protocols utilizing T4 RNA ligase, adapter ligation is influenced by the ligase used, the miRNA insert and adapter primary sequence, as well as the GC content and the secondary structures of miRNA insert and adapter^19–23^. For poly-A utilizing protocols, the enzyme poly (A) polymerase has also been reported to be influenced by miRNA primary sequence and secondary structure^24^ Other reported possible sources of bias during library preparation include the reverse transcription and PCR steps, with PCR in particular able to introduce both amplification bias and duplicate reads, but results have been contradictory^19–21^. A recent study recommended the use of unique molecular identifiers (UMI) to mitigate the reverse-transcription and PCR biases in future experiments^25^. Previous studies also reported that the incorporation of UMIs into sequence adapters resulted in improved accuracy both in RNA-seq and smallRNA-seq analysis^26 27^

In this study we aimed to systematically assess the miRNA repertoire and frequency observed in NGS data using six different low-input library preparation protocols (Table 1). Commercial vendors marketing kits stating compatibility with total RNA amounts ≤100 ng were invited to participate. The performance of the protocols was compared with regard to their detection rate sensitivity, reliability and ability to identify differentially expressed miRNAs. In addition, the relevance of UMIs was studied. All analyses were performed on low-input well-defined synthetic miRNA and human-derived total RNA samples.

**Table 1:** Small RNA library preparation methods tested in this study.

| **Method Name** | **Commercial supplier** | **Key points\*** | **Max. input volume tolerated** | **Reported RNA input range (varies with type of input tested)** | **Maximum number of indexes available** | **Method types** |
| --- | --- | --- | --- | --- | --- | --- |
| CATS Small RNA-Seq Kit (CATS) | Diagenode s.a. | Single-tube, 4-step process of polynucleotide tailing, reverse transcription and PCR amplification. 1 purification step (2 if size selection required). | 8 µl | 0.1 – 100 ng | 24 | Poly-A based. 2-adapter procedure |
| Small RNA-Seq Library Prep Kit (srLp) | Lexogen GmbH [Cat. No. 052, 058] | 4-step procedure of 3' adapter ligation, 5' adapter ligation, reverse transcription and PCR. 2 purification steps (3 if size selection required). | 6 µl | 0.05 – 1000 ng | 96 | Ligase based. 2-adapter procedure |
| QIAseq miRNA Library Kit (QIAseq) | QIAGEN [Cat. No. 331502 or 331505] | 5-step procedure of 3' adapter ligation, 5' adapter ligation, reverse transcription and PCR. 2 purification steps (3 if size selection required). | 5 µl | 1 – 500 ng | 96 | Ligase based. UMI incorporated. 2-adapter procedure |
| TailorMix microRNA Sample Preparation Kit Version 3 (TailorMix) | SeqMatic LLC. | 4-step procedure of 3' adapter ligation, 5' adapter ligation, reverse transcription and PCR. 2 purification steps, including a final PAGE gel excision. | 6 µl | 1– 1000 ng | 96 | Ligase based. 2-adapter procedure |
| SMARTer® miRNA-seq Kit (Beta version) (SMARTer) | Takara Bio USA Inc. | 5-step procedure of 3' mono-adapter ligation, dephosphorylation, adapter dimer blocking, circularisation, reverse transcription and PCR. 2 purification steps, including a final PAGE gel excision. | 4 µl | 100 – 1000 ng total RNA or 2 – 200 ng enriched small RNA | 48 | Ligase based. Single adapter procedure |
| CleanTag™<br>Small RNA<br>Library Prep<br>Kit<br>(CleanTag) | TriLink<br>BioTechnologi<br>es, LLC. | Single-tube, 4-step<br>procedure of 3' adapter<br>ligation, 5' adapter ligation,<br>reverse transcription and<br>PCR. 1 purification step. | 10µl | 1 – 1000 ng | 48 | Ligase<br>based.<br>2-adapter<br>procedur<br>e |

\* A step is defined as a labwork period that culminates in an incubation longer than 5 minutes.

## Results

### Experimental design and miRNA read yields

Synthetic miRNA and biologically derived human total RNA samples (21 samples in total) were distributed to participating companies for library preparation (Figure 1a). Upon return of libraries, library yield and size were measured (Supplementary Figure 2). Libraries were pooled based on their miRNA content with the aim of obtaining at least 5 million reads apiece. Based on the presence of additional small RNAs in some libraries, particularly those prepared with the CATS and SMARTer-beta reagents, which displayed a wider range of insert sizes, we did not expect all libraries to produce similar read counts at this stage. Nevertheless, a large number of reads from both CATS and SMARTer-beta kits failed to pass QC threshold filters.

**Figure 1.**
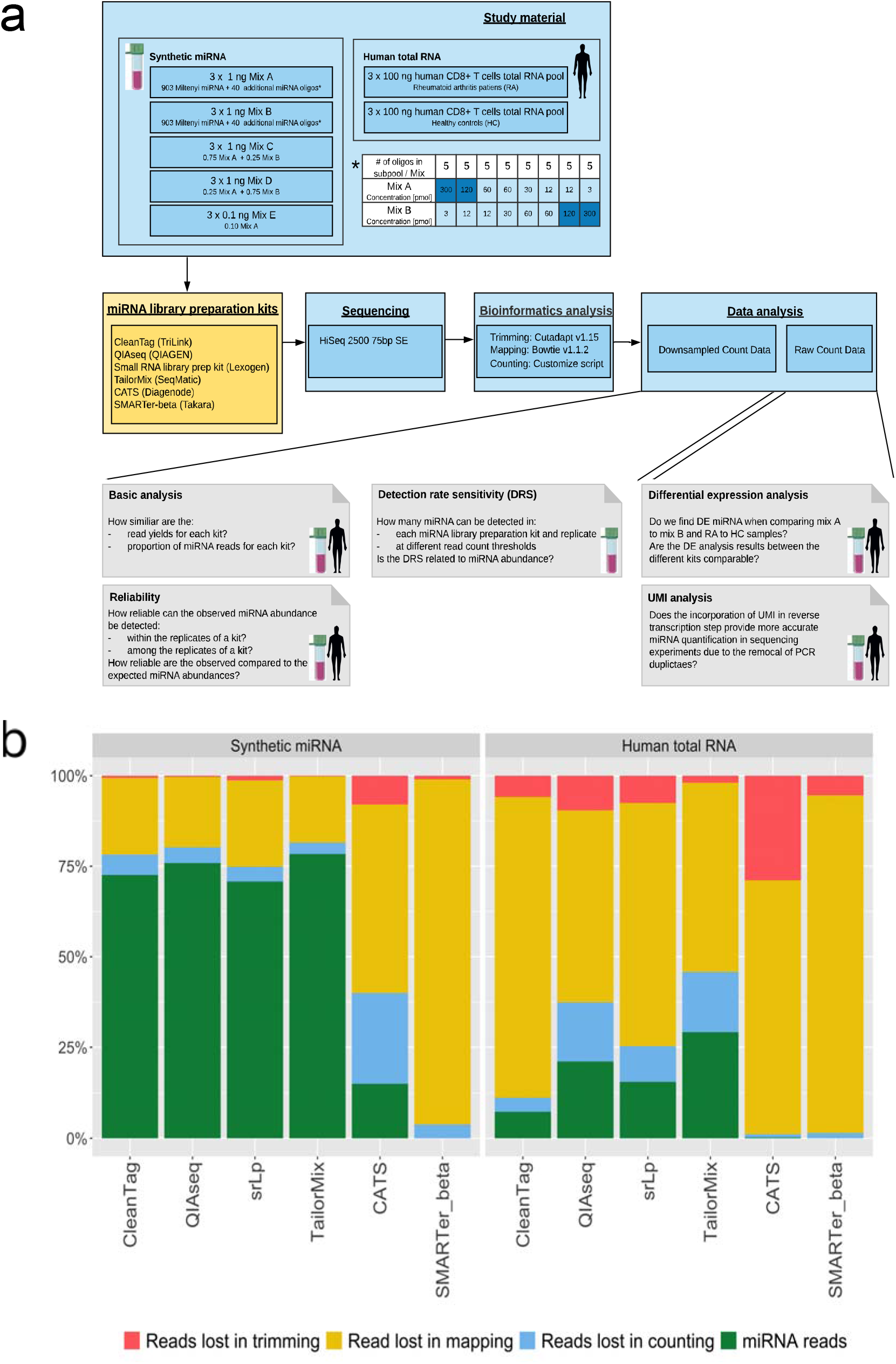
Experimental design and sequencing read distribution. A: Overview of the study material, miRNA library preparation kits used, sequencing, bioinformatics and data analysis. Steps presented in blue boxes were performed in-house, while the step presented in the yellow box was executed by the indicated library preparation vendors. Grey boxes represent individual data analysis steps. B: Percentage of reads that were removed during the bioinformatic analysis and final miRNA proportion remaining (green). Trimming refers to removal of adapter sequences, mapping to miRNA reference alignment, and counting to filtering of aligned miRNAs that did not have the same length as the reference sequence. Results presented are the mean of 15 replicates in the synthetic miRNA (left) and the mean of six replicates in the human total RNA samples (right). Figure 1 A was created using images from Servier Medical Art (Servier. www.servier.com, licensed under a Creative Commons Attribution 3.0 Unported License).

The number of sequencing reads obtained ranged from 400,000 to more than 33 million reads for the individual libraries (Supplementary Table 4).

For all library preparation kits, the greatest proportion of reads were discarded during mapping, most likely as a result of not allowing for any mismatches (Figure 1b and Supplementary Figure 3). The absolute number of reads excluded differed between the kits. As anticipated, a higher fraction of miRNA reads was obtained in the synthetic miRNA samples compared to the human total RNA samples (Figure 1b), since the human total RNA samples also contain additional classes of small RNA. SMARTer-beta and CATS returned the lowest proportion of miRNA reads both in the synthetic miRNA and the human total RNA samples compared to the other library preparation kits while TailorMix, followed by QIAseq produced the highest proportions of miRNA reads (Figure 1b).

To comprehensively evaluate the sensitivity and reliability of the library preparation kits, the synthetic miRNA samples were randomly down-sampled to 2.5 million and human total RNA samples to 0.75 million miRNA reads. The libraries of SMARTer-beta and CATS did not reach these thresholds and were therefore excluded from further analysis. The results presented hereafter are based on calculations using the down-sampled data with the exception of the differential expression and UMI analyses for which raw (not down-sampled) miRNA reads were used.

### Detection rate sensitivity

Applying a relaxed detection threshold where miRNAs were defined as detected if one or more read counts were registered, the detection rate sensitivity for all kits ranged from 94.7% to 99.1%, and all miRNAs could be detected in at least one kit and replicate. QIAseq followed by TailorMix detected the highest numbers of miRNAs in all three replicates in all the mixes (Figure 2a). QIAseq and TailorMix also missed the fewest miRNAs in either one, two, or all three triplicates. When comparing the detection rate sensitivity of the 1.0ng synthetic miRNA samples (mix A-D) with the 0.1ng synthetic miRNA samples (mix E), no striking difference in the number of detected miRNAs could be observed for any of the kits (Figure 2a).

**Figure 2.**
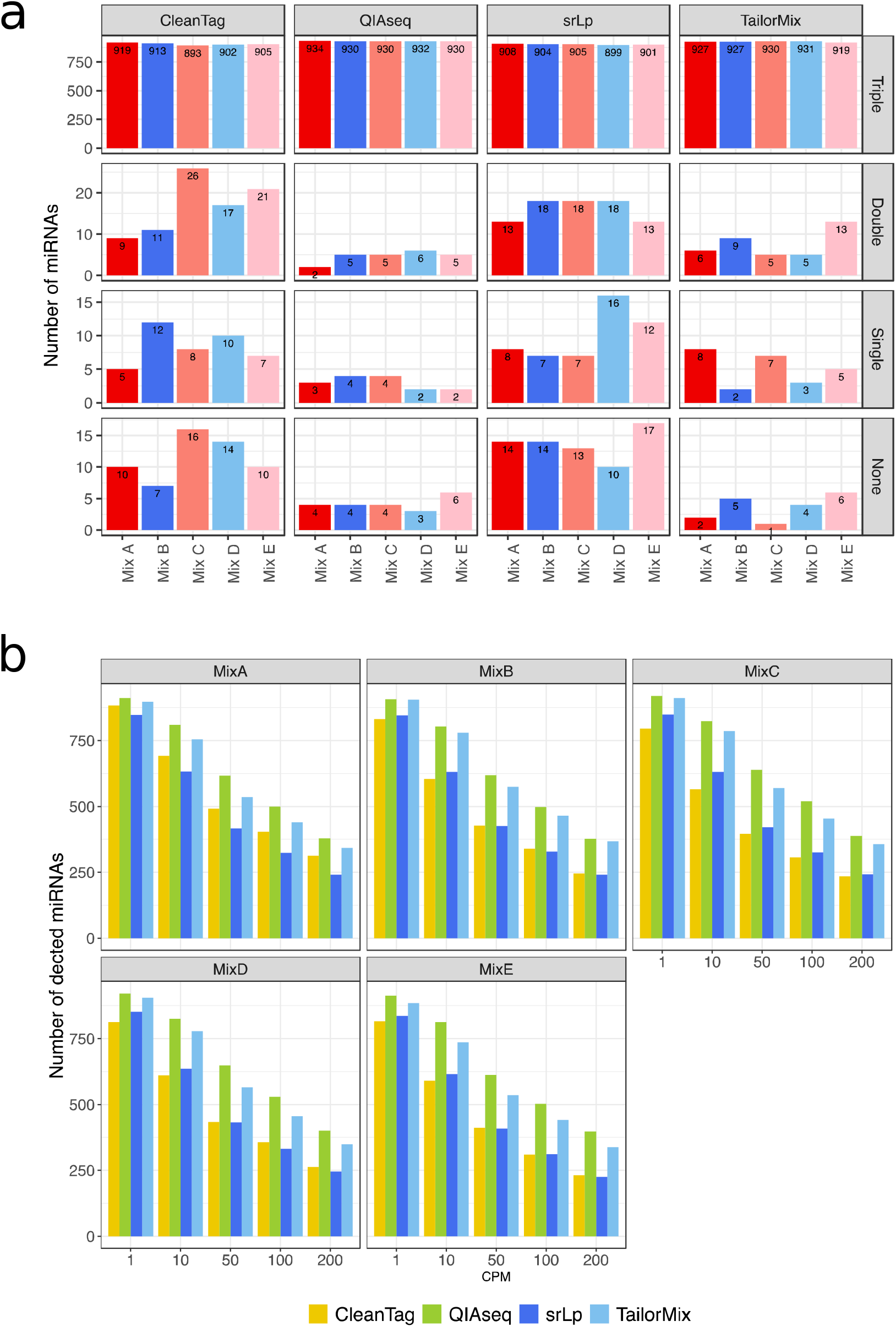
Detection rate sensitivity. A: Bar charts presenting number of miRNAs detected in all replicates (Triple), in 2 out of 3 replicates (Double), in 1 out of 3 replicates (Single) or not detected in any replicate (None) across all synthetic miRNA mixes and all library preparation kits. The maximum number of detectable miRNAs is 943 (903 equimolar and 40 non-equimolar miRNA). B: Bar charts for various read count thresholds in the synthetic miRNA samples. A miRNA is defined as detected when it is (i) expressed in all three replicates of the mix and (ii) the read counts are greater or equal to the count per million (CPM) threshold displayed on the x-axis. The colours of the bars represent the reagents.

Most of the miRNAs that were undetected in QIAseq and TailorMix were neither detected by the other two kits (Supplementary Figures 4 and 5). TailorMix was the only reagent that detected each of the 903 equimolar miRNAs in at least one sample (Supplementary Figure 5). srLp followed by CleanTag showed the highest numbers of kit-specific undetectable miRNAs.

Analysis of the 40 non-equimolar miRNAs revealed that miRNAs undetected in one or more replicates belonged mostly to miRNAs present at low levels (Supplementary Figure 6), with QIAseq showing the highest detection rate, again followed by TailorMix. Notably, CleanTag and srLp failed to detect some miRNAs present at relatively high concentrations in all the replicates (mix C and D, Supplementary Figure 6). However, even though the majority of the non-equimolar miRNAs could be detected in all replicates, the analysis indicated that factors in addition to miRNA abundance influence detection rate sensitivity.

We next compared the performance at different detection thresholds, i.e. 1, 10, 50, 100, 200 read counts per million (CPM) for synthetic miRNA samples in all mix triplicates for each kit (Figure 2b). With the exception of some of the non-equimolar miRNA oligonucleotides present at the lowest concentration, all synthetic miRNAs should in theory obtain CPM values above 200 with the library size of 2.5 million mapped miRNA reads. However, a sharp decline in detection was observed at increasing CPM thresholds. Nonetheless, QIAseq followed by TailorMix consistently detected the highest number of miRNAs across all thresholds.

### Intra-rater and Inter-rater reliability

Rlog transformed miRNA count data were used for the for intra- and inter-rater reliability calculations. Intra-rater reliability calculations (the concordance between miRNA read counts within the replicates of the library preparation kit) revealed excellent reliability for the synthetic miRNA and the human total RNA samples within all tested kits with ICC values above 0.99 and 0.98, respectively (Supplementary Table 5). Similarly, very strong correlations were found when Pearson correlation coefficients were calculated (r > 0.97, p < 0.05) (Supplementary Figure 7 and 9a). Bland-Altman plots, which describe the agreement between two replicates by presenting the difference of them against the mean, also showed good agreement (Supplementary Figure 8 and 9b). For all Bland-Altman comparisons the bias was close to 0. The line of equality (not presented in our Bland-Altman plots) was always within the agreement limits, which indicates a lack of systematic error in the measurements within the replicates. All in all, strong intra-rater reliabilities were observed within the samples prepared by each kit.

For the inter-rater reliability calculations (concordance of read counts seen between the different kits) the first replicate of each mix, RA or healthy control sample was randomly chosen. The synthetic miRNA and the human total RNA samples revealed good and excellent inter-rater reliability with ICC values above 0.83 and 0.95 respectively (Supplementary Table 6). The correlation between the different kits was above 0.76 (p < 0.05) for the synthetic miRNA and above 0.92 for the human total RNA samples (Supplementary Figure 10). However, differences in the correlations between the reagents were seen for the synthetic miRNA samples. The kits with the highest correlations (r > 0.94, p < 0.05) were, independent of whether mix A-E was considered, CleanTag and srLp while QIAseq showed the lowest correlation to the other kits. The Bland-Altman plots revealed no systematic error when comparing the different kits to each other (Supplementary Figures 11 and 12). The limits of agreements were smallest for CleanTag and srLp across all tested mixes in the synthetic miRNA samples indicating a high agreement between those two kits. In summary, a modest-to-good inter-rater reliability was obtained when comparing the mix-specific replicates of the four miRNA library preparation kits with each other, with QIAseq showing the greatest differences from the other reagents.

The reliability measured against the theoretical miRNA concentration was only assessed for the synthetic miRNA samples. For the 903 equimolar miRNAs, the fold deviation of the first replicate of mix A from the median count for that sample was calculated as a rlog ratio (Supplementary Figure 13). When the absolute value of the rlog fold deviation for a miRNA was less than or equal to one, the miRNA was counted as equimolar. For the four kits this was the case for 39.8 to 42.0% of the equimolar miRNAs. The remaining miRNAs showed a bias towards over-representation (positive rlog fold change) rather than under-representation. The coefficient of variation of the rlog counts across all replicates for the equimolar miRNAs was lowest for QIAseq, followed by TailorMix, CleanTag and srLp, respectively (Supplementary Table 7).

For the 40 non-equimolar miRNAs, the correlation between the rlog counts of each library preparation kit and their theoretical concentration varied between the mixes for all kits. Overall, mix A and mix E showed greater correlations (0.41 < r > 0.61, p < 0.05) than mix B to mix D (0.08 < r > 0.47, p < 0.05) (Supplementary Table 8). QIAseq showed the highest correlation coefficients across all samples. All in all, these results suggest that on one hand the reliability between the reagents is good, but on the other hand that none of the reagents are ideally suited for accurate miRNA quantification.

### Differential expression

Most miRNA profiling studies aim to identify differentially expressed (DE) miRNAs between samples of interest. When comparing mix A and mix B of the synthetic miRNA samples, ideally all 40 non-equimolar miRNAs should be detected as DE with a log2 fold change greater than or equal to one. All kits detected between 32 to 35 DE miRNAs (Figure 3a). However, some of those miRNAs (CleanTag, TailorMix and srLp=2 and QIAseq=1) were from the pool of equimolar miRNAs. Of the 40 non-equimolar miRNAs, 26 were detected to be DE by all kits, although they did not always agree on the log fold changes (Figure 3b). The non-equimolar miRNAs hsa-miR-1199-5p, hsa-miR-22-5p and hsa-miR-940, which were three of the ten miRNAs expected to show the lowest fold differences (fold change of 2) between mix A and mix B, could not be detected as DE by any of the reagents.

**Figure 3.**
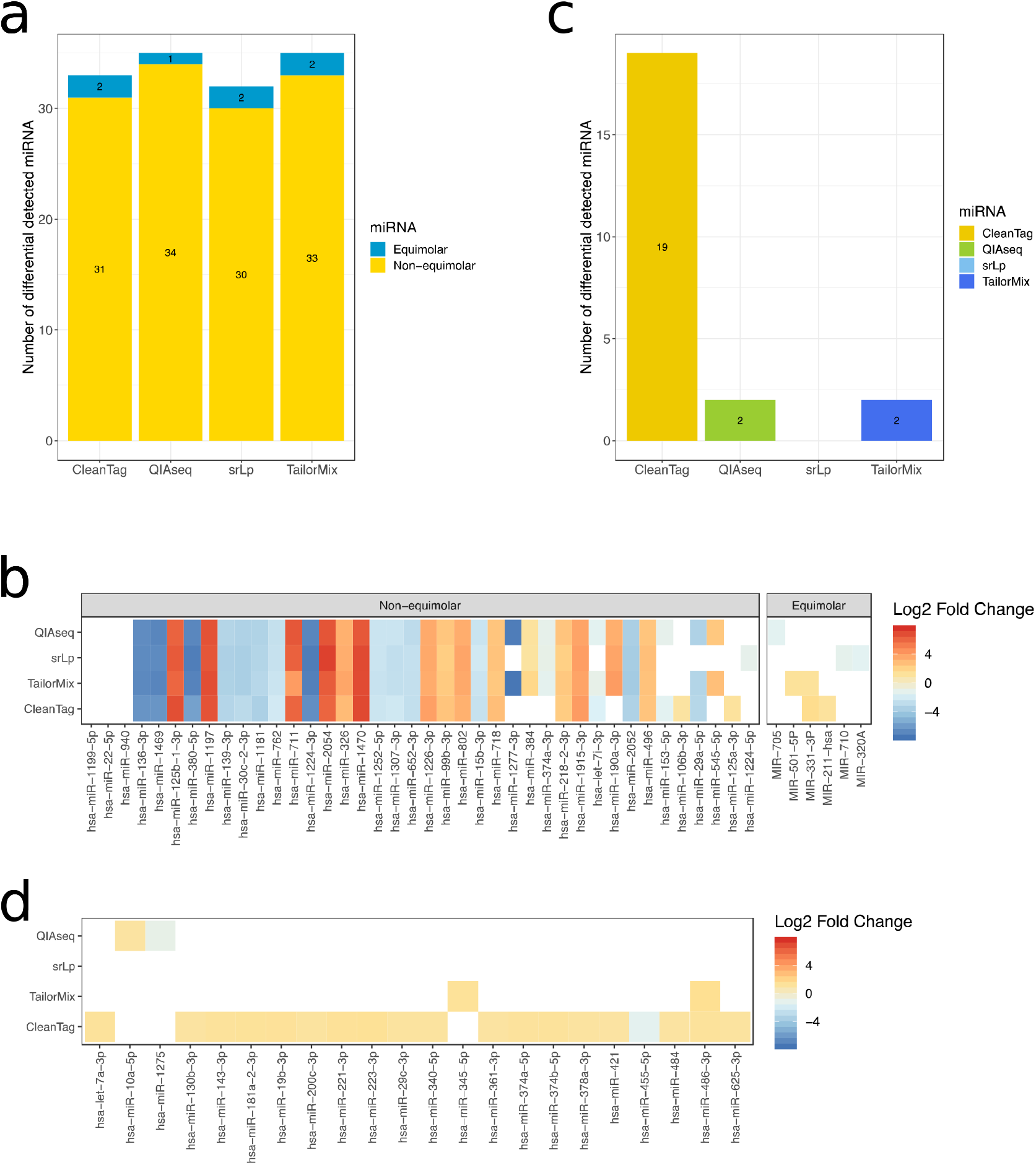
Differential expression analysis. Kit-specific number of differentially expressed miRNA detected for A: synthetic miRNA samples (mix A versus mix B) and C: human total RNA samples (RA versus healthy control). miRNA-specific log2 fold changes across the different kits for B: synthetic miRNA samples and D: human total RNA samples.

In order to control as best possible that the levels of miRNA in mix A and mix B were as expected, we performed quantitative reverse-transcriptase PCR assays on 16 selected nonequimolar miRNAs (Supplementary Figure 14), which confirmed the intended ratios in the starting material.

Differential expression analysis of the human total RNA samples revealed different numbers of DE miRNAs detected by the kits. CleanTag detected 19 DE miRNAs, QIAseq and TailorMix detected two DE miRNAs each, while srLp did not detect any (Figure 3c). With the exception of hsa-miR-486-3p, no overlap between the DE miRNAs was seen amongst the kits (Figure 3d).

### Titration response

The titration response of the 40 non-equimolar miRNAs in mixes A – D (Figure 1a) was compared by scoring a miRNA as titrating or non-titrating based on detection in the expected concentration order in the four mixes. Since there were five miRNAs at each chosen concentration, the fraction of titrating miRNAs (0, 0.2, 0.4, 0.6, 0.8 or 1) was calculated for each reagent kit for each concentration group (Table 2). The highest fraction of titrating miRNAs was seen for QIAseq, which correctly scored all miRNA concentrations with greater than 2-fold differences in mix A through mix D.

**Table 2:**
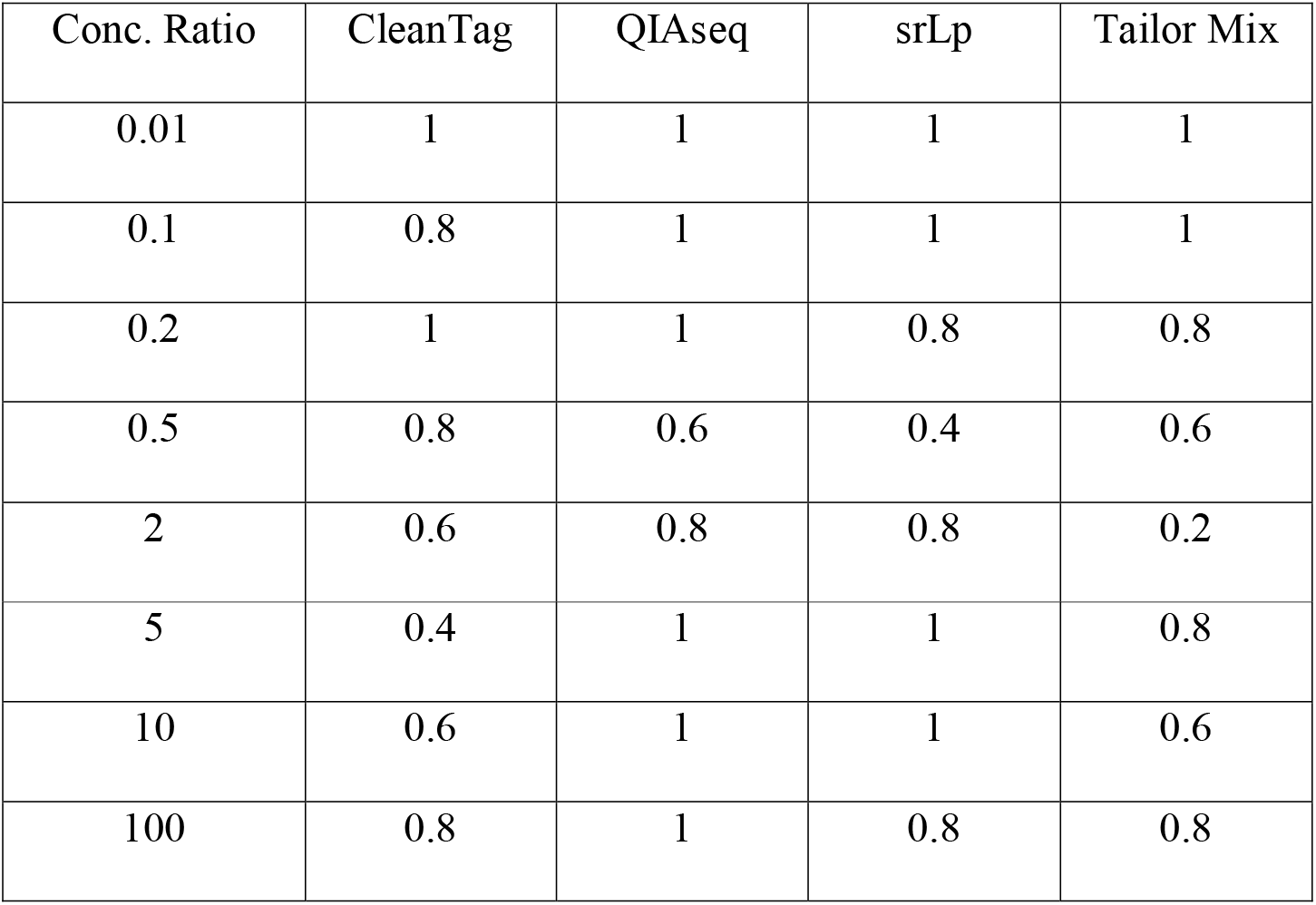
Fraction of titrating miRNAs (n=5) in each of the eight concentration groups. Average rlog expression values for the 40 non-equimolar miRNAs were calculated across the three replicates each of mixes A to D. Each miRNA was scored as titrating if the average values followed the expected trend in concentrations from high to low or vice versa across mixes A to D.

| Conc. Ratio | CleanTag | QIAseq | srLp | Tailor Mix |
| --- | --- | --- | --- | --- |
| 0.01 | 1 | 1 | 1 | 1 |
| 0.1 | 0.8 | 1 | 1 | 1 |
| 0.2 | 1 | 1 | 0.8 | 0.8 |
| 0.5 | 0.8 | 0.6 | 0.4 | 0.6 |
| 2 | 0.6 | 0.8 | 0.8 | 0.2 |
| 5 | 0.4 | 1 | 1 | 0.8 |
| 10 | 0.6 | 1 | 1 | 0.6 |
| 100 | 0.8 | 1 | 0.8 | 0.8 |

### Effectiveness of QIAseq unique molecular identifier sequence tags

QIAseq was the only kit included in this study that implements unique molecular identifiers (UMIs) during library preparation, which are claimed to enable more accurate quantification of miRNAs. For both synthetic miRNA and human total RNA samples, very strong Pearson correlations were observed between the rlog transformed raw read and UMI counts (Supplementary Figure 15). Comparison of the rlog sum of all UMI and ordinary read counts revealed the sum of UMI counts to be negligibly smaller than the ordinary read counts for both synthetic miRNA and human total RNA samples (Supplementary Figure 16a, b).

To further examine whether UMI read counts might reduce undesirable over-representation of miRNAs that were favourably amplified or sequenced, we examined the abundance of the ten miRNAs with the highest ordinary read counts for each sample and compared this to their respective UMI counts (Supplementary Figure 16c,d). Amongst those miRNAs no overestimation of the ordinary read counts was observed compared to the UMI counts.

## Discussion

Several publications have revealed discrepancies between the frequencies of miRNAs present in the original samples and those detected by sequencing approaches^18 20^. The adapter ligation steps in the small library preparation procedure, in addition to miRNA sequence and structure, have emerged as being most critical when trying to explain the discrepancy^18 19 21 43 44 20 22^ As an alternative to the ligase-dependent ligation step in library preparation, polyadenylation based procedures have been developed. Additional biases might be introduced during reverse transcription and PCR steps, but in this case results have been contradictory (^19–21, 25^). The use of UMI tags has therefore been suggested to remove this potential bias^25^. Here we performed a comprehensive comparison of six low input small RNA sequencing reagents utilizing both ligase-depend, polyA-based and single-adapter methods, including one kit that employed UMI tags. Note that we assessed here only the performance of the kits to identify miRNAs; other small RNA species that may be captured were not assessed.

### Sequencing yields and miRNA read proportions

Considerably different numbers of raw reads were obtained from the different kits. The kits from TailorMix and QIAseq returned the highest miRNA read counts both in the synthetic miRNA and the human total RNA samples. However, raw read outputs cannot be used to judge the performance of a method. Furthermore, since the samples from SMARTer-beta were sequenced alone in a single lane, we cannot exclude that technical issues affecting only that lane were responsible for the low raw read numbers that passed filters. The input range tested in this study was at or below the indicated range stated for the SMARTer-beta kit (100 ng −1 ug total RNA or 2 ng – 200 ng enriched small RNA); this may have resulted in the observed poor performance. Since this study was performed, the kit has been re-optimized and released with a new formulation and improved performance. Nonetheless, the low proportion of reads mapping to miRNAs from both the CATS and SMARTer-beta was clearly evident, which could be attributable to inefficient removal of other small RNA species during library preparation. However, greater numbers of reads that were not counted as miRNA (due to imperfect match in length to the database reference sequence) were noticeable for CATS, which may indicate that polyadenylation-based methods are trickier to process during data analysis, due to uncertainties on the length of the poly-A tail added. To reduce the influence of technical aspects (e.g. different library size selection and purification methods, as well as raw read yields) on the comparisons, all miRNA counts were downsampled to the same levels. CATS and SMARTer-beta did not reach the selected thresholds and were therefore excluded from further analysis.

### Detection rate sensitivity

When applying low detection rate sensitivity thresholds, most synthetic miRNAs could be detected by the four remaining kits, indicating that all of them may be suited to assess the overall miRNA repertoire. However, when applying more stringent detection thresholds ranging from 1cpm to 200cpm, greater differences in detection rates between the kits became evident, and QIAseq and TailorMix emerged as the most sensitive. It is worth noting that kit specific biases played a greater role in miRNA detection than input RNA amounts, at least within the ranges tested here (0.1-1.0 ng miRNA).

### Reliability

Intra-rater reliability showed very high concordance between miRNA counts within the replicates of a miRNA library preparation kit, independent of the kit, for both synthetic miRNA and human total RNA inputs. Similar results have been reported by Giraldez, et al.^44^ and Wright, et al.^25^, although they refer to intra-rater reliability as reproducibility and consistency respectively. The intra-rater reliability was strong both for 0.1ng and 1.0ng synthetic miRNA samples for all kits in our study (data not shown) which is promising given current interest in using low RNA inputs derived from small biological specimens.

In concordance with the findings reported by Giraldez, et al.^44^, Coenen-Stass, et al.^45^ and Wright, et al.^25^, inter-rater reliability (concordance of read counts seen between the different kits, also called reproducibility or consistency across replicates) was lower compared to the intra-rater reliability. In particular, QIAseq deviated from the other kits, but we stress that this does not indicate poorer performance. QIAseq employs a different 3’adapter sequence compared to the other three kits which may underlie the dissimilar preference for subset of miRNAs observed. These observations underscore the emerging conclusion that kit-specific differences should be considered by any researchers comparing miRNA-seq datasets, as supported by another recent study^46^ Notably, the concordance between the miRNA counts measured and the expected concentration for the synthetic miRNA samples was low, and revealed that none of the library preparation kits could accurately quantify the majority of miRNAs.

### Differential expression

Differential expression analysis of synthetic miRNA mix A versus mix B revealed that all kits could detect at least 31 out of 40 non-equimolar miRNA correctly as DE (fold change ≥ 2). MiRNAs hsa-miR-1199-5p, hsa-miR-22-5p and hsa-miR-940 were never detected as DE by any of the kits. These miRNAs were present at two-fold concentration differences, the lowest fold change tested, which can be challenging. In general, all reagents displayed greater problems to detect small fold-change differences, reminiscent of results seen in the recent study by Giraldez, et al.^44^

Our study offered the additional possibility to study levels of false positive DE miRNAs detected from the 903 equimolar miRNAs. Equimolar miRNAs found to be DE were characteristically detected as DE with low fold-changes and showed little agreement between the kits, consistent with their being false positive calls. Taken together, QIAseq showed slightly higher sensitivity (true positives) and slightly higher specificity (fewer false positives) than the other reagents, although the false-positive calls did fall within the expected rate set for the analysis (False discovery rate = 0.05). Reinforcing these conclusions, the titration response analysis clearly demonstrated the superior performance of the QIAseq reagents to most faithfully represent the levels of miRNAs in input material.

It nonetheless appears that the different reagents have differing preferences for particular miRNAs. The primary sequence of terminal miRNA nucleotides^18^, secondary structure affecting ligation sites^47^ and co-folding of the miRNA and ligated adapters^21^ have all been documented as sources of bias affecting miRNA detection. Interestingly, the 3’ adapter sequence in the QIAseq kit differs from the other three kits analysed. However, our attempts to explain the differences observed between the kits based on primary sequence or secondary structure analyses were inconclusive (data not shown).

Greater differences between kits were observed by examining DE miRNAs detected when comparing the RA patient pool and healthy control pool of human CD8+ T cell RNA, where the number of DE miRNA varied between none (srLp) to 19 (CleanTag). There are few preceding studies of miRNAs from blood-isolated CD8+ cells in rheumatoid arthritis, but some of the miRNAs found to be DE in this study have previously been associated with RA, e.g. miR-221-3p^48^, miR-223-3p^49–51^, miR-374b-5p^52^ and miR-486-3p^52^, however further confirmation is needed. Worryingly, in addition to the varying number of DE miRNA detected by the different kits, there was almost no concordance between the miRNAs identified. Taken together, it is advisable to interpret DE miRNA results from studies employing different library preparation methods with caution.

### Re-analysis of QIAseq dataset utilising UMIs

Reverse transcription and PCR-amplification may be potential sources of bias during library preparation, and PCR can also introduce duplicate reads. QIAseq was the only kit tested to address the issue of duplicate reads by the inclusion of UMIs, however, under the employed conditions, no appreciable difference between UMI counts and the ordinary read counts were detected, mirroring the findings of Wong, et al.^46^. Fu, et al.^27^ observed that higher fractions of PCR duplicates could be observed when reducing the starting material, but when comparing the 1.0ng and 0.1ng synthetic miRNA samples, no difference in the proportion of PCR duplicates was seen. Nonetheless, it remains possible that at lower concentrations than tested here, UMIs may prove useful for the elimination of duplicates to improve dataset quality.

In conclusion, the QIAseq kit from QIAGEN consistently demonstrated performance at, or near, the top for all metrics examined. It should be mentioned that QIAGEN made an error affecting samples 1-8 in their first attempt at library preparation and were supplied with replacements. With the exception of performance in the titration response assay, the TailorMix kit from SeqMatic closely followed. Lexogen’s srLp and Trilink’s CleanTag kit also performed well, and the majority of differences we detected point to kit-specific biases. However, whilst the experiments conducted here show that sequencing is a very sensitive method for detecting miRNAs, even at low abundance, it is also clear that none of the kits performed impressively with regard to accurately reflecting the relative input levels of all miRNAs. There is clearly room for improvements in this regard for the development of further enhanced reagents or methods to accurately quantitate miRNA levels.

## Material and Methods

### Study material

The performance of six miRNA library preparation kits was examined using low-input material consisting of synthetic miRNA samples or human-derived total RNA samples. To maximize the possibility that each procedure was performed under optimum conditions, samples were distributed to the kit vendors for library construction. Sequencing libraries were returned to the Norwegian Sequencing Centre for sequencing and data analysis.

### Synthetic miRNA samples

The synthetic miRNA samples consisted of a mixture of equimolar and non-equimolar miRNAs. The miRXplore Universal Reference (Miltenyi, California, United States), comprising 962 HPLC purified, 5’ phosphorylated, synthetic oligonucleotides of human, mouse, rat and viral miRNA origin, was used as an equimolar miRNA pool. For the non-equimolar pool, 40 additional HPLC purified, 5’phosphorylated, synthetic oligonucleotides representing human miRNA were purchased from Eurofins MWG Synthesis GmbH (Bavaria, Germany). Altogether five different miRNA mixes were created (denoted mix A to mix E, Figure 1a). Mix A and Mix B consisted of the equimolar miRNA pool supplemented with the non-equimolar pool present at eight different concentration ratios between the two mixes spanning a 100-fold range (Supplementary Table S1). Mix C was a titration of 0.75 mix A and 0.25 mix B, while mix D was a titration of 0.25 mix A and 0.75 mix B. In the case of mixes A-D, the total miRNA concentration was 30 nM, with individual equimolar miRNAs present at 30 pM and other miRNAs ranging from 3 – 300 pM. Mix E consisted of the same miRNAs as mix A but at a 10-fold lower concentration. Due to the low concentrations in the five synthetic miRNA mixes, the samples were blended with yeast (*Saccharomyces cerevisiae*) total RNA, which does not contain known endogenous miRNAs^28^, to minimise degradation and loss of material due to adhesion to plasticware, and to mimic the more complex total RNA mixtures encountered under typical usage. In each mix, the final RNA content was 2 ng/μl, with miRNA representing approx. 10% (w/w) of the total amount (mixes A-D) or 1% (mix E). The samples were distributed in triplicates to the participating vendors. To each of the triplicates in mix A to mix E, one additional specific miRNA (miR-147a, miR-212-3p or miR-412-3p) was added to check that the replicates were processed independently throughout library preparation and were not combined into a single sample to increase reproducibility.

To verify the intended ratios of the synthetic miRNA sample starting material, quantitative reverse-transcriptase PCR was performed using 16 pre-designed TaqMan^®^ Small RNA assays (Thermo Fisher Scientific, Waltham, MA USA) according to manufacturer’s instructions. Assay details are provided in Supplementary Material and Methods. Relative abundances of miRNAs in mixes A and B were measured by absolute quantification relative to a standard curve.

### Human-derived total RNA samples

Peripheral blood CD8+ T cells were magnetically sorted from newly diagnosed rheumatoid arthritis (RA) patients (n=4) and healthy controls (n=4) using the EasySep cell isolation system (Stemcell technologies, Vancouver, Canada). The RNA/DNA/Protein Purification Kit (Norgen Biotek, Ontario, Canada) was used to isolate total RNA. Only RNA samples with RNA integrity values above 8.5 were used for downstream analysis. To ensure the desired amount of total RNA input for the miRNA library preparation, the four RA patients and the four healthy controls were mixed together to obtain one pooled RA and one pooled healthy control sample respectively. Triplicates of these different sample types were distributed to the participants.

### miRNA library preparation

Each participant was asked to prepare miRNA libraries from the 21 samples described above using their specific miRNA library preparation kit. For optimization purposes the participants received a further 20 ng of synthetic miRNA (blend of Mix A and Mix B) and 200 ng total human RNA. All participants were requested to use the same Illumina i7 index sequence for the same sample to avoid any possible effect of these sequences on the downstream library preparation and sequencing process. Detailed sample and index information can be found in Supplementary Table S2.

At the time of writing, four of the six kits were commercially available in the formats used for this study (CATs, QIAseq, CleanTag and TailorMix). A fifth kit, srLp, was also commercially available, but with different index primer sequences. For comparison purposes and to avoid possible bias arising from the use of different indexes, this participant synthesised custom index primers complying with the index sequences specified in this article. The SMARTer kit used in the study had not been released for purchase, but a modified version is now available. It should be noted that this study is not exhaustive, since two library preparation suppliers meeting the input amount inclusion criteria (PerkinElmer, formerly Bioo Scientific, and NEB) declined to participate. Detailed descriptions of the library preparation conditions employed by the producers of the specific reagents are supplied in the Supplementary Material and Methods.

### Sequencing

All libraries were sequenced at the Norwegian Sequencing Center on the same single-read flow cell of a HiSeq 2500 (Illumina, San Diego, CA) with 75 bp reads generated using v4 clustering and SBS reagents according to the manufacturer’s instructions. To avoid sequencing lane bias, the libraries of srLp, QIAseq, TailorMix, CATS and CleanTag were randomly distributed over five lanes of the flow cell, equivalent to sequencing 21 libraries per lane (Supplementary Table S4). Due to concerns that the SMARTer beta libraries contained a large proportion of non-miRNA inserts (higher molecular weight products than expected, making it challenging to obtain equivalent numbers of reads per sample), these libraries were sequenced independently from the other participants on a single lane (Supplementary Figure 1),

## Bioinformatic analysis

### Read mapping and reference sequences

Primary base calling and quality scoring was performed using RTA v1.18.66.4 (Illumina), followed by demultiplexing and processing with Bcl2fastq v2.18.0.12 (Illumina). For trimming of the 3’ adapter, cutadapt v1.15^29^ with parameter –m 10 was used. Detailed information about adapter sequences is provided in the Supplementary Material and Methods. Read mapping was performed using bowtie v1.1.2^30^ with parameters –a and --norc. No mismatch was allowed. As reference, the expected pools of synthetic miRNAs (962 synthetic equimolar miRNAs originating from the miRXplore universal reference and 40 non-equimolar miRNAs) were used for the synthetic miRNA samples, and the mature human miRNA sequences specified in miRBase^31^ v21 for the human total RNA samples. We confirmed that all replicates had been processed separately by verifying the presence/absence of spiked replicate-specific miRNAs in the datasets from each sample. Further analysis revealed that 59 of the miRNA sequences included in the miRXplore Universal Reference were identical to sequences in the *Saccharomyces cerevisae* (sacCer3) genome (Supplementary Table S3). To avoid potential miscounting of yeast fragments in the downstream analysis, these miRNA were excluded and only the remaining 903 miRNA of the miRXplore Universal Reference were analysed further. Mapped reads (restricted to miRNAs matching exactly to the reference sequence and length) were counted using a custom python script (available upon request).

### Read count modelling

With the exception of differential expression and UMI analysis, all further downstream analyses were performed on down-sampled mapped miRNA reads to minimise confounding factors arising from sources such as read numbers and proportions of adapter dimer reads, which can be influenced by the purification method chosen and by pipetting errors. Random down-sampling to 2.5 million reads was performed for the synthetic miRNA samples and to 0.75 million reads for the human total RNA samples. The seed number was set to 123.

In miRNA-seq count data, the average observed variance across samples increases with higher average expression of the miRNA. If this heteroscedastic behaviour of the count data is not taken into account, the results of most downstream analyses will be dominated by highly expressed and highly variable miRNAs. We therefore transformed count data, where indicated, with the rlog function of DeSeq2^32^ (v1.20.0), which produces a superior homoscedastic output than log2 transformation for low- and high-expressed genes^32^

## Data analysis

### Detection rate sensitivity and reliability

Data and statistical analyses were performed using R v3.5.2^33^ and Python v2.7.13. Unless otherwise stated, ggplot2^34^ was used for data visualization. Synthetic miRNA and human total RNA down-sampled read count data were used in the detection rate sensitivity analysis. Upset plots were produced using the R package UpSetR^35^ v1.4.0.

Rlog transformed synthetic miRNA and human total RNA count data were used for assessing the reliability of the library preparation kits, on which intra-class correlation (ICC), Pearson correlation and Bland-Altman agreements calculations were performed. For ICC, the two-way mixed effects model, absolute agreement and single rater (ICC(3,1)) were applied using the R package psych^36^ v1.8.4. ICC values were interpreted according to the recommendations of Koo and Li^37^ where ICC values above 0.9, between 0.75 and 0.9, between 0.5 and 0.75 and below 0.5 indicate excellent, good, moderate and poor reliability respectively. Thresholds described by Chan^38^ were used for the Pearson correlation where correlations above 0.8, between 0.6 and 0.8, between 0.3 and 0.6 and below 0.3 are described as very strong, moderately strong, fair and poor respectively. The R corrplot package^39^ v0.84 was utilized for correlation plots and the R BlandAltmanLeh package^40^ v0.31 for Bland Altman calculations.

### Differential expression and titration response

Original read count data of mix A and mix B were used for the differential expression analysis using the R package edgeR^41^ v3.22.3. For the synthetic miRNA samples a read count filtering of 3 counts per million (cpm) in at least two libraries was applied to the differential expression analysis while a filter of 20cpm in at least two libraries was used for the human total RNA samples. miRNAs were defined as significantly differentially expressed after multiple testing adjustment with the methods of Benjamini and Hochberg controlling for a false discovery rate of 0.05. In addition, only those miRNA with |log2 FC| > 1 between the tested conditions were kept.

The titration response of the 40 non-equimolar miRNAs of the synthetic miRNA samples was examined in mixes A to D according to the analyses published by Shippy, et al.^42^ Average rlog expression values for each miRNA were calculated across the three replicates of each of mixes A to D. If the average expression values for each miRNA followed the expected concentration trend (across the four possible concentrations seen in each mix), it was scored as titrating. Any deviations from the expected trend were scored as non-titrating.

### UMI analysis

QIAGEN’s analysis tool Geneglobe was used for assessing the effectiveness of QIAseq’s UMIs. For the synthetic miRNA samples the option “other” was chosen for mapping while “human” was chosen for the human total RNA samples during the primary data analysis. The resulting count table included UMI (after PCR duplicate removal) and raw (before PCR duplicate removal) read counts for each miRNA in the samples. Before analysing the correlation between UMI and raw read counts, the counts were rlog transformed.

## Acknowledgements

We thank Iris Langstein and Philipp Korber for *S. cerevisiae* RNA. Dalia Daujotyte and Stephanie Bannister are also thanked for reviewing the results and manuscript.

## Data availability statement

Raw sequencing fastq files and miRNA count tables will be made available in the Gene Expression Omnibus database.

## Appendices

**Supplementary Figures** (document: Supplementary_figures_Heinicke_etal2019.docx)

**Supplementary Tables** (document: Supplementary_tables_Heinicke_etal2019.xlsx)

**Supplementary Material and Methods** (document: Supplementary_Material_and_Methods_Heinicke_etal2019.docx)

