## Supplementary Figures for "Systematic assessment of commercially available low-input miRNA library preparation kits"

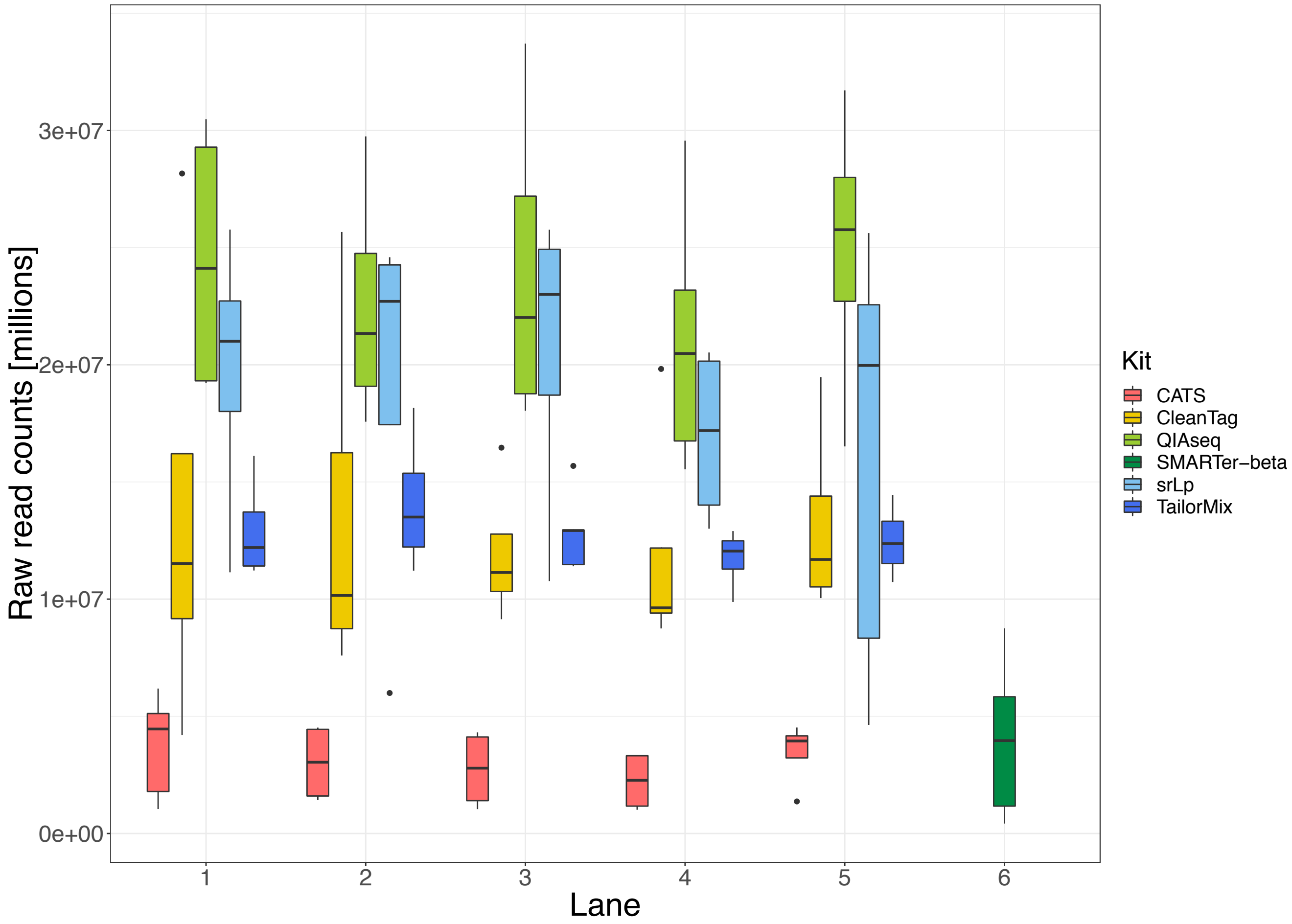


**Figure S1: Kit-specific raw read distributions obtained on six lanes of a HiSeq2500 flow cell. Based on their miRNA library contents, indexed libraries (with the exception of SMARTer-beta) were pooled across lanes 1-5 to avoid any flow cell lane bias. Indexed SMARTer-beta libraries were sequenced on a separate lane of the flow cell.**

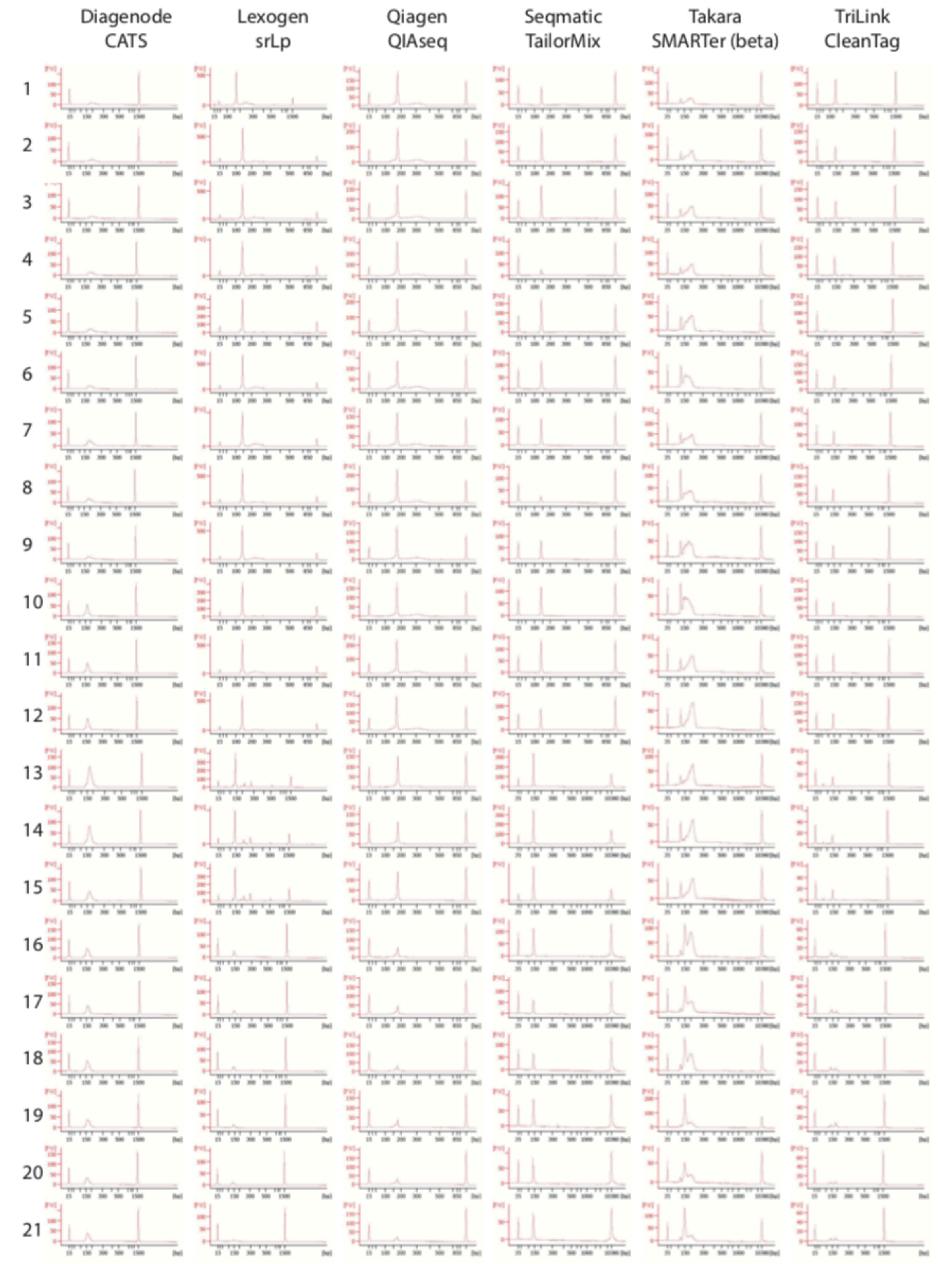


**Figure S2: Bioanalyzer electrophoretograms of all participant libraries returned for sequencing. X-axis in all cases is size in bp, and Y-axis fluorescence intensity (nucleic acid amount). Peaks at 15bp and 1500bp are internal reference size markers.**

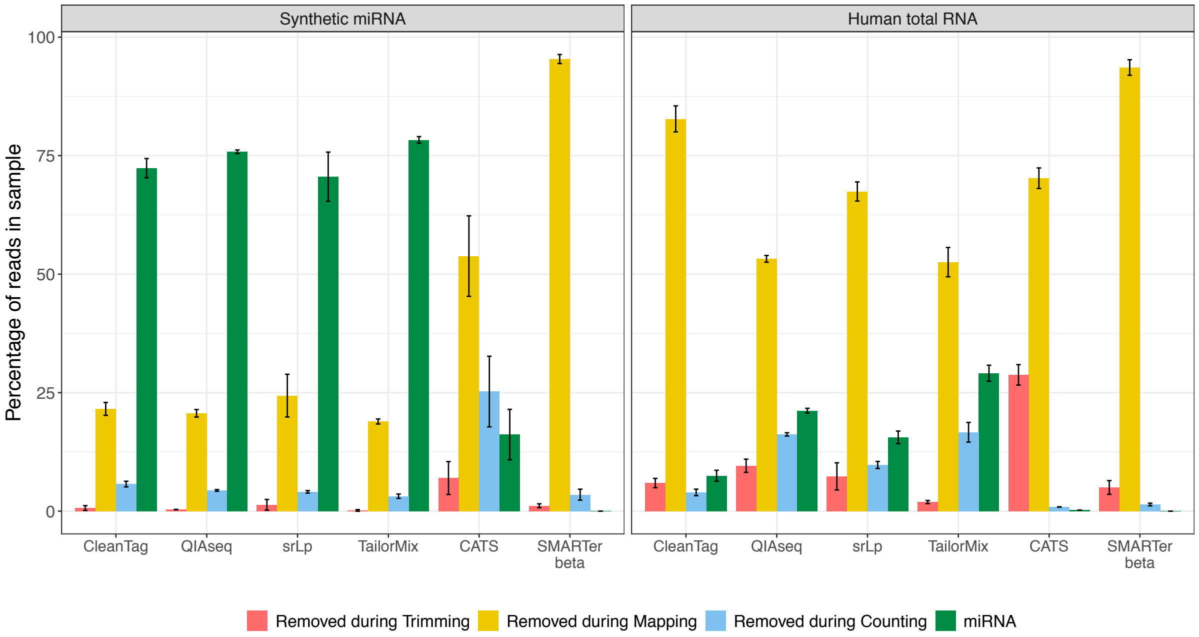


**Figure S3: Sequencing read distribution. Percentage of reads that were removed during the bioinformatics analysis and remaining miRNA reads. The results presented are the mean of 15 replicates in the synthetic miRNA and the mean of six replicates in the human total RNA samples. Error bars represent mean read count +/- standard deviation.**

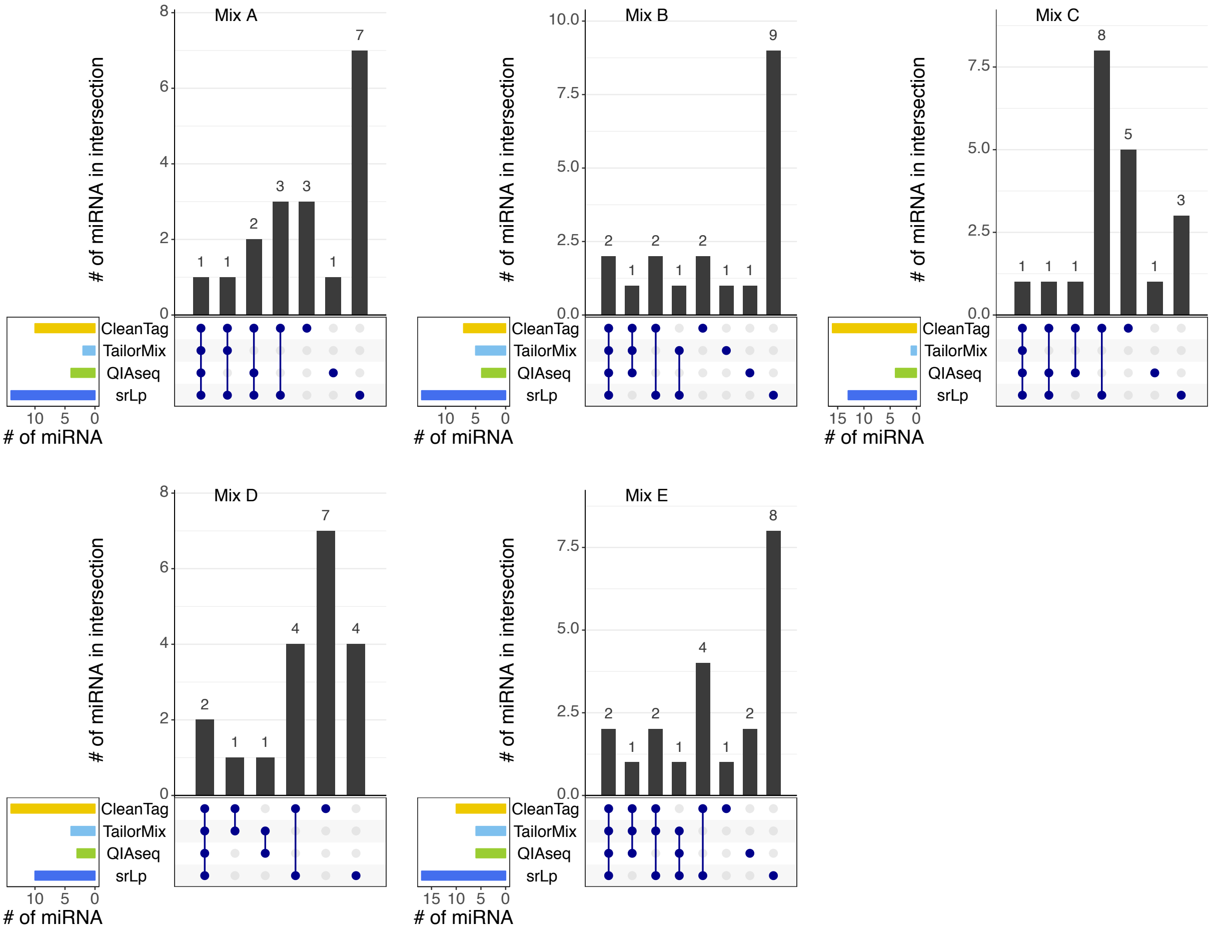


**Figure S4: Upset plots representing the intersections of undetected synthetic miRNAs across the different kits and mixes. The interactions are indicated by the blue dots and lines. The number of miRNAs in the intersection is represented by the black bar charts.**

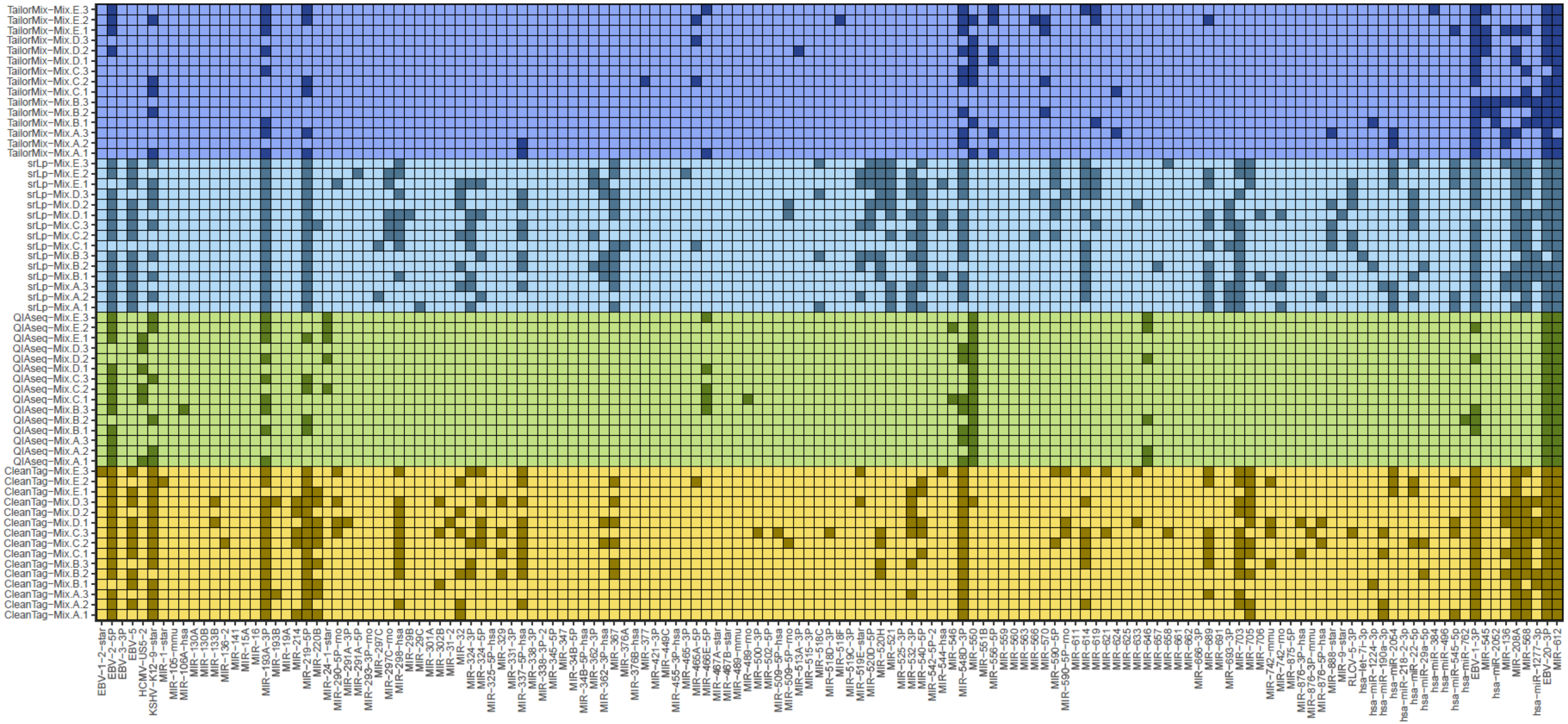


**Figure S5: Detection rate sensitivity for the synthetic miRNAs that could not be detected in at least one of the replicates for one of the kits and in at least one mix. The remaining miRNAs not represented in the plot were detected in all replicates of all kits. The library preparation kit and the replicate are presented on the y-axis. A darkened box within the plot indicates that the miRNA for this kit and replicate was not detected with at least one count.**

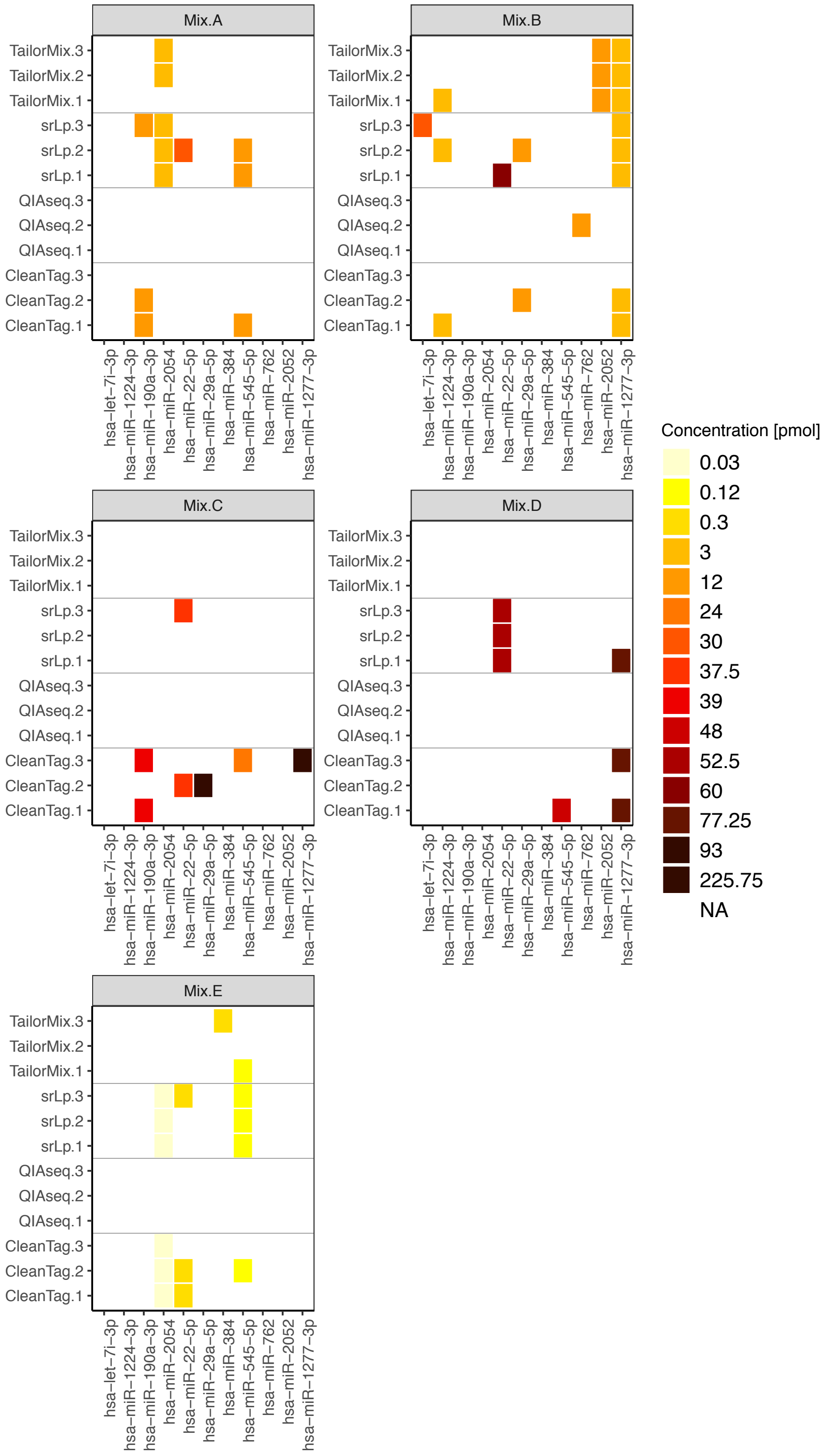


**Figure S6: Detection rate sensitivity. Each of the five subplots present the detection rate sensitivity for the non-equimolar miRNAs in mix A to mix E. miRNAs presented on the x axis (n=11) represent miRNAs that could not be detected in at least one of the replicates of one kit and in one mix. The remaining 27 miRNAs of the non-equimolar group (not presented) were detected in all replicates of all reagents in all five mixes. The library preparation reagents and the replicate are presented on the y-axis. A coloured box within a plot indicates that the miRNA for this kit and replicate could not be detected. The colour of the box represents the concentration of the miRNA.**

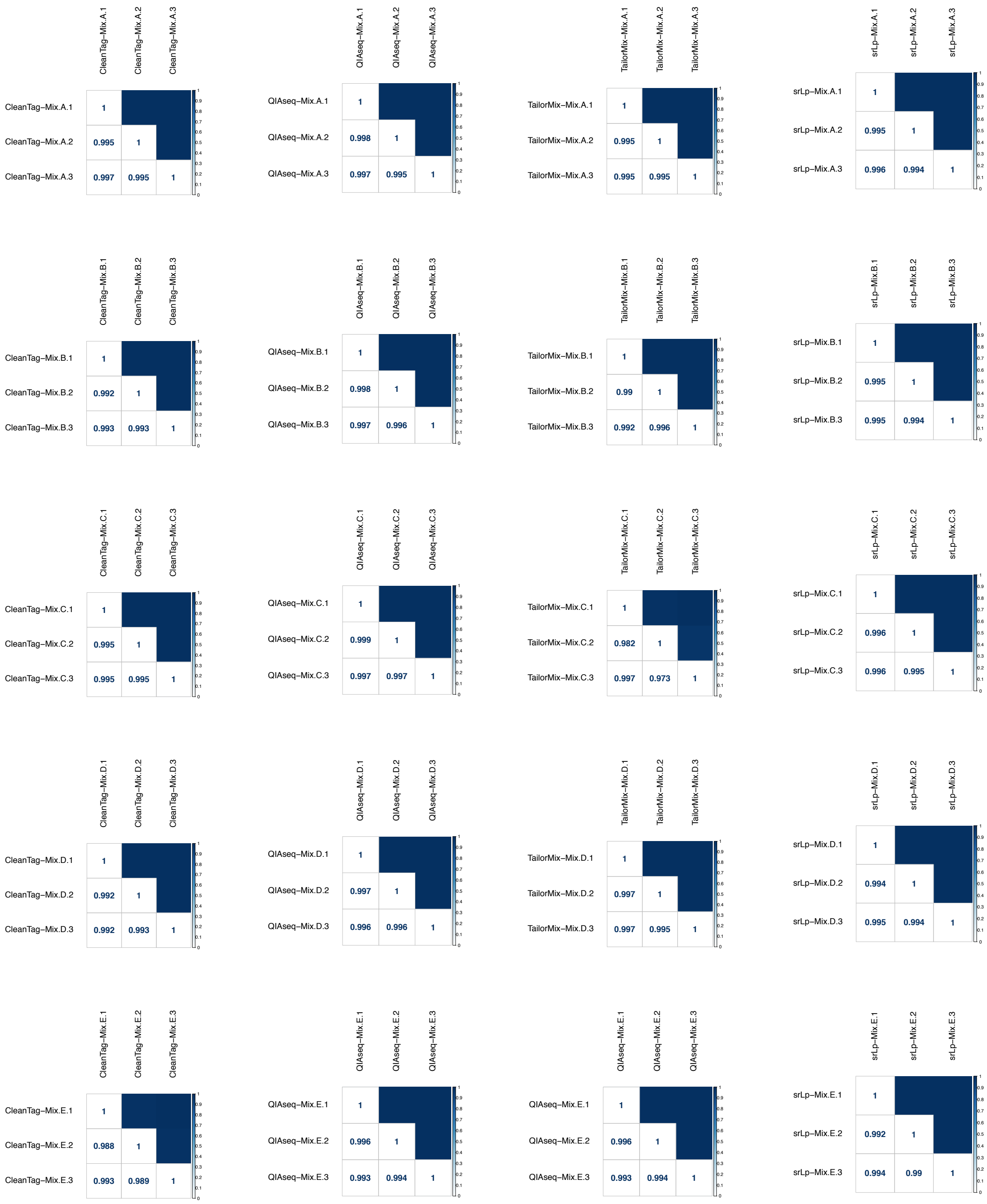

Figure S7: Intra-rater Pearson correlation coefficients of the read counts within the replicates of the kits for the synthetic miRNA samples. Each correlation square presents a specific replicate, mix and kit.

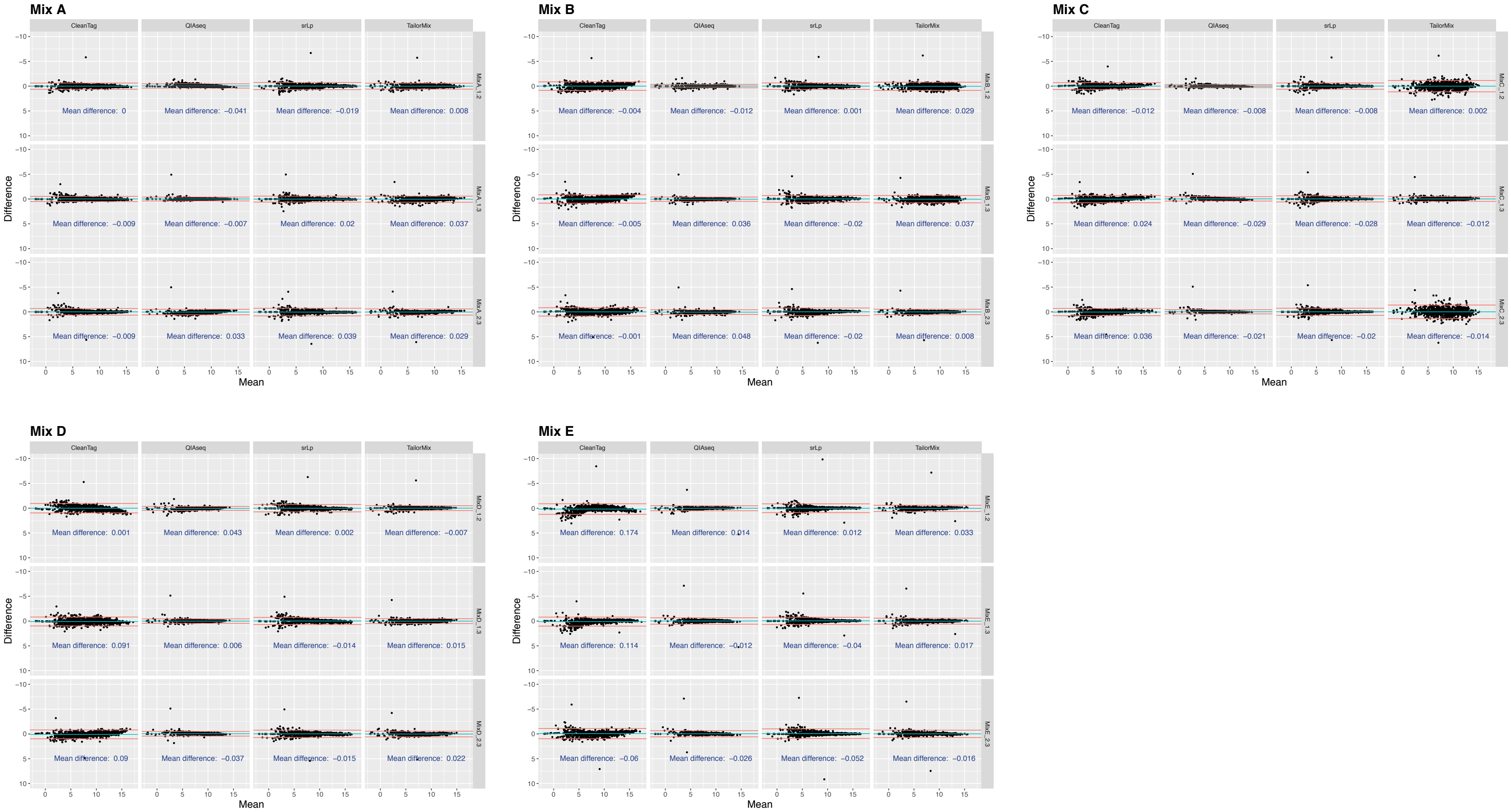


**Figure S8: Bland-Altman plots of the intra-rater analysis showing the differences against the average of two specific replicates within a kit for all synthetic miRNA oligos in mix A to mix E. The black dots represent the miRNAs while the blue line and the two red lines represent the bias (also called mean difference and the 95% CI agreement limits respectively).**

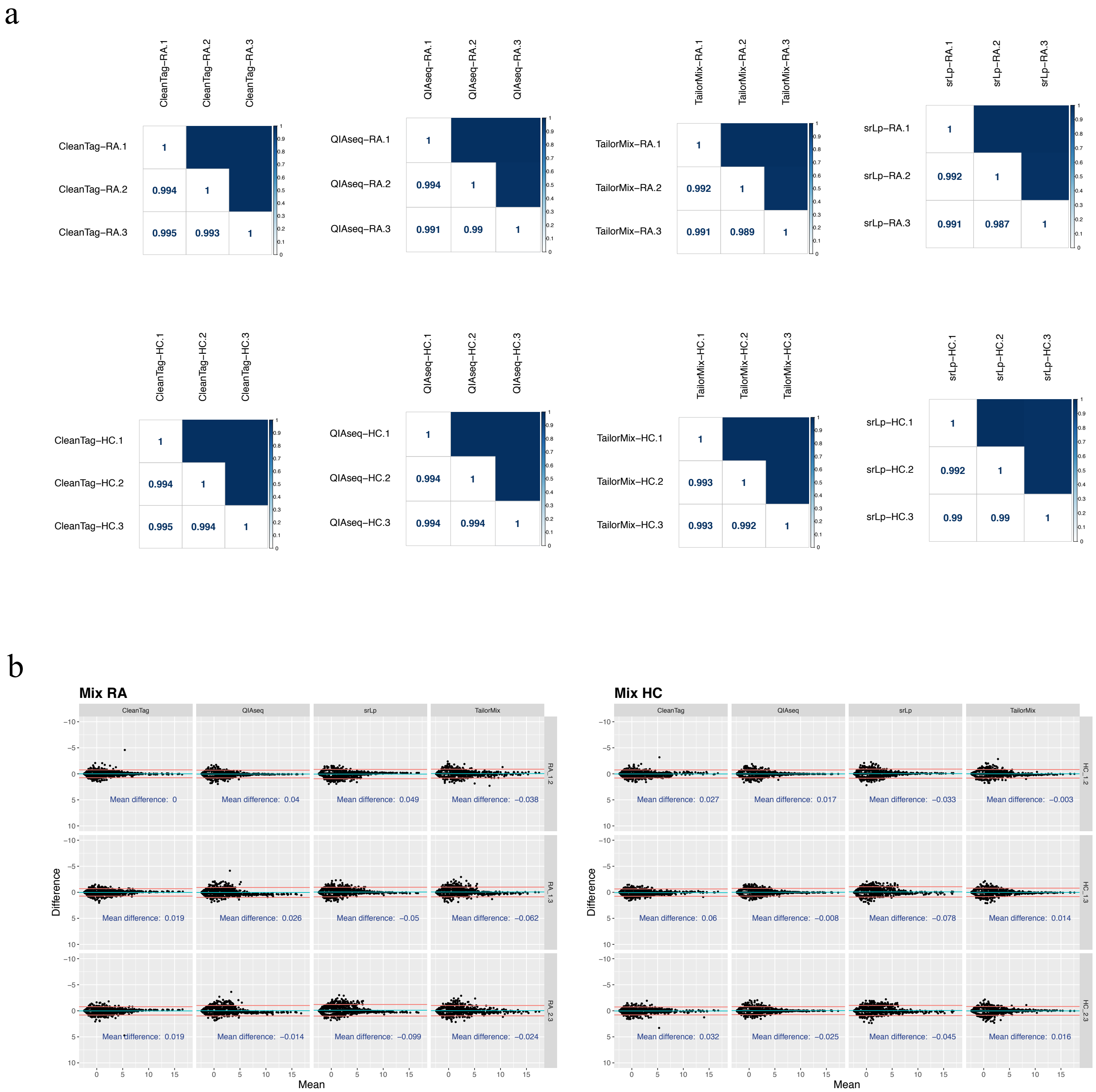


**Figure S9: Intra-rater reliability analysis in the human total RNA samples. (a) Pearson correlation of the read counts within the replicates of the various kits for rheumatoid arthritis (RA) and healthy control (HC) samples. Each correlation square presents a specific replicate, mix and kit. (b) Bland-Altman plots show the differences against the average of two specific replicates within a kit type. The black dots represent individual miRNAs while the blue line and the two red lines represent the bias (also called mean difference and the 95% CI agreement limits).**

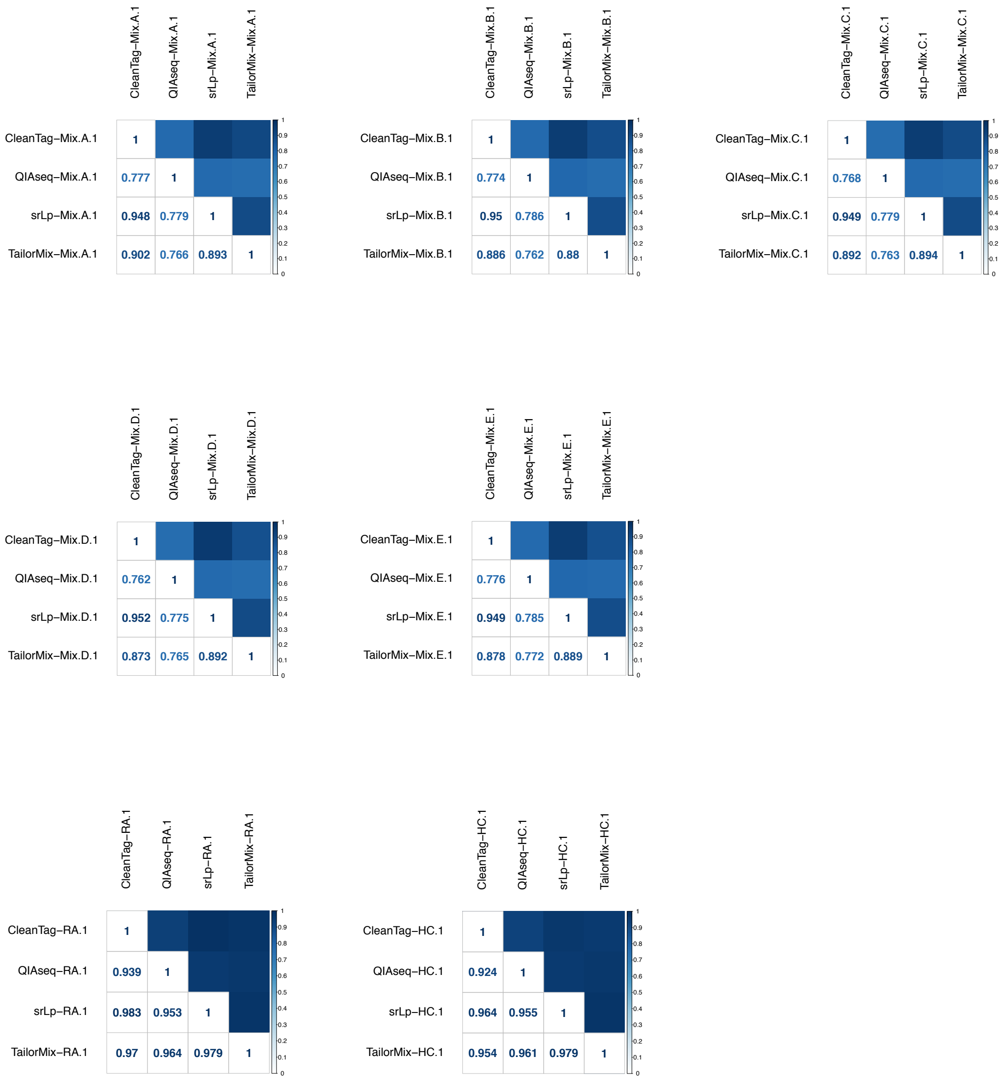


**Figure S10: Inter-rater correlation of the read counts within the first replicate of all five mixes in each kit for the synthetic miRNAs, rheumatoid arthritis (RA) and healthy control (HC) samples. Each correlation square presents a specific mix and kit.**

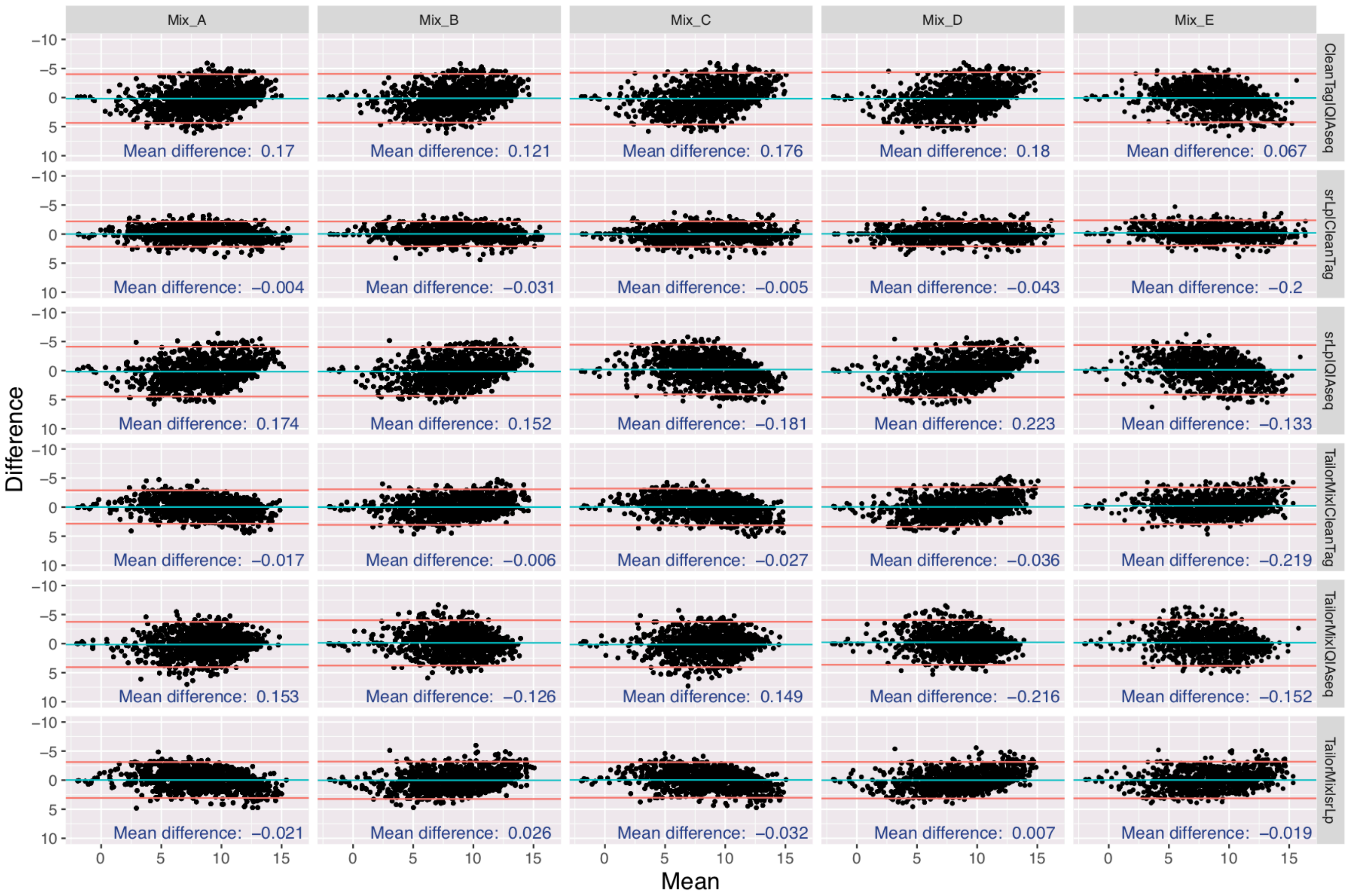


**Figure S11: Bland-Altman plots of the inter-rater reliability in the synthetic miRNA samples for the first replicate of all five mixes in each kit. Bland-Altman plots show the differences against the average of two specific kits within mix A to mix E. The black dots represent individual miRNAs while the blue line and the two red lines represent the bias (also called mean difference and the 95% CI agreement limits).**

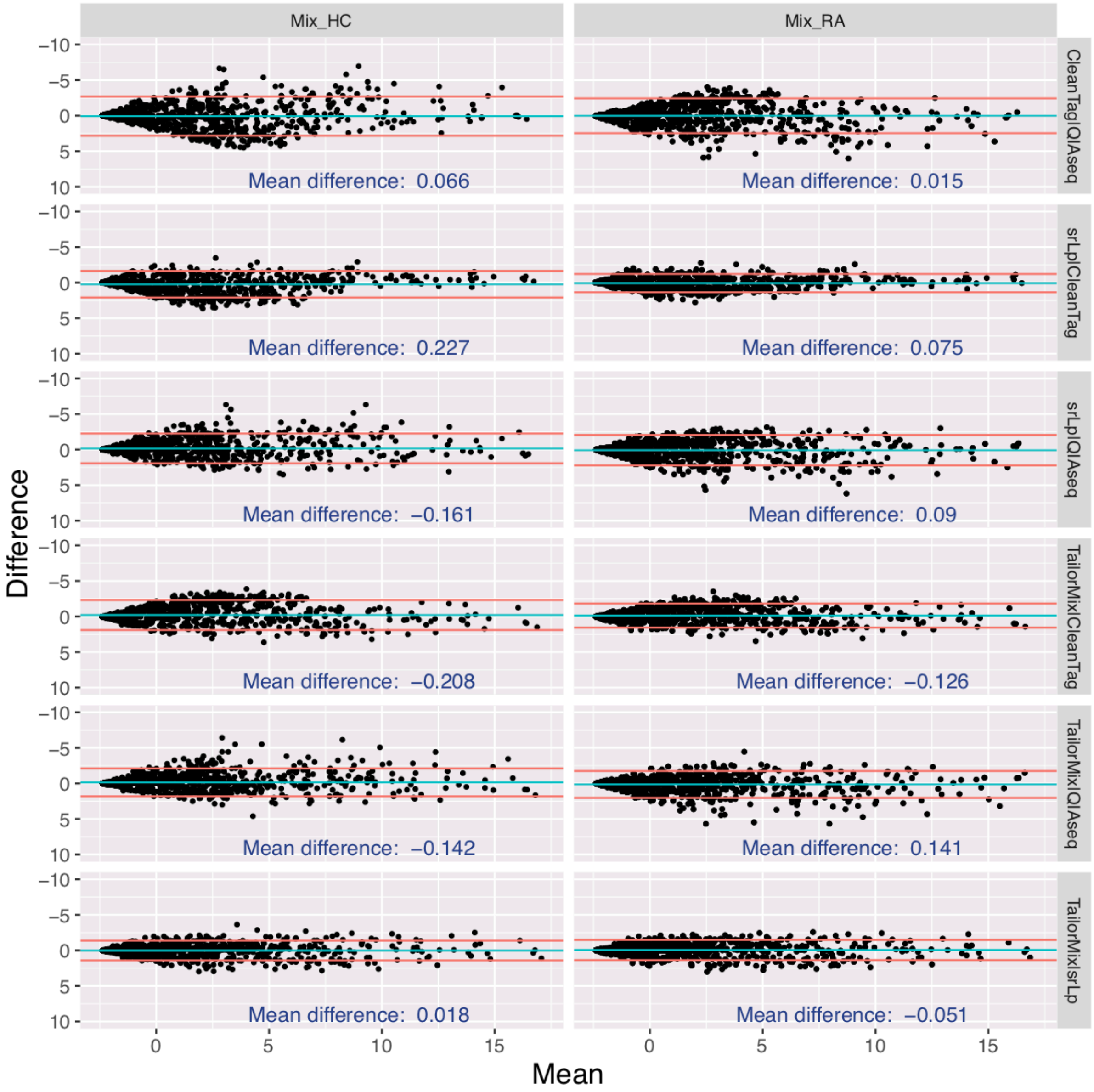


**Figure S12: Bland-Altman plots of the inter-rater reliability for the first replicate of rheumatoid arthritis (RA) or healthy control (HC) samples in each kit. Bland-Altman plots show the differences against the average of two specific kits. The black dots represent individual miRNAs while the blue line and the two red lines represent the bias (also called mean difference and the 95% CI agreement limits).**

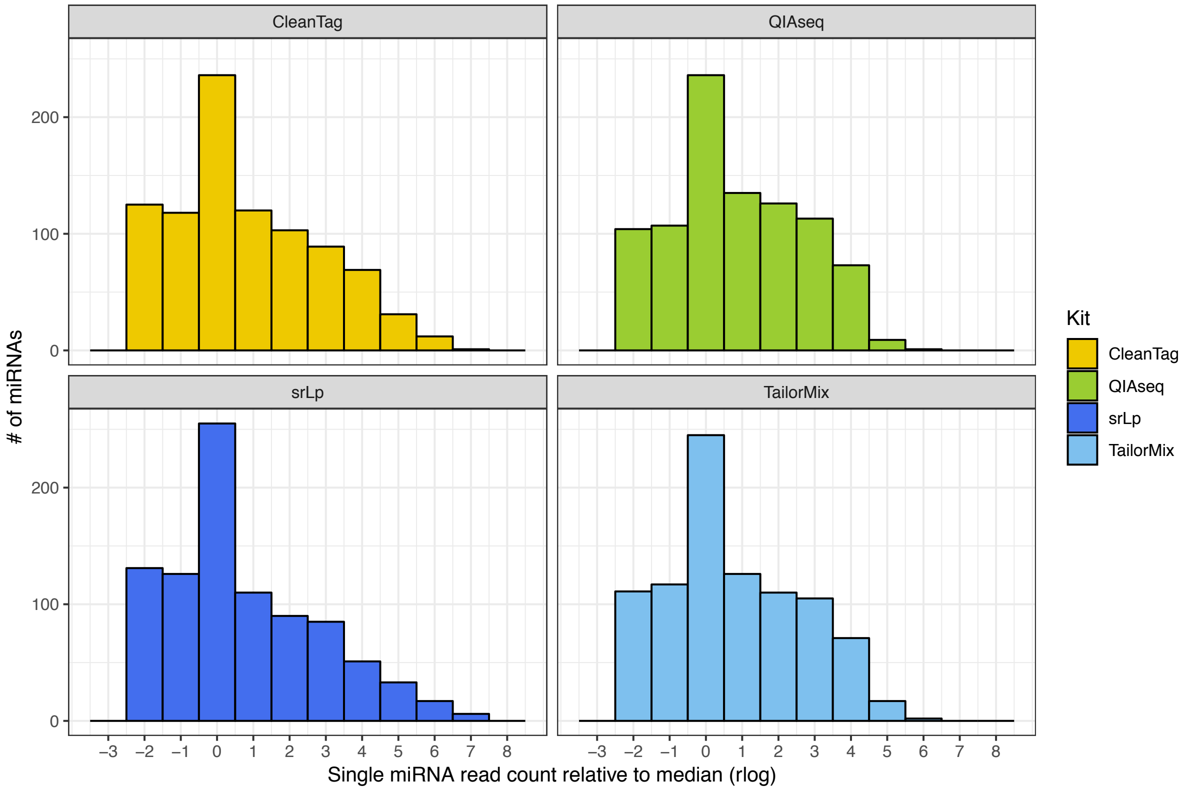


**Figure S13: Bias in miRNA detection utilizing the synthetic equimolar miRNA reference miRXplore. Bar charts represent the rlog ratio of single miRNA read counts to the median read counts of all the equimolar miRNAs of the specific library preparation kit. Results for replicate 1 of synthetic mix A are shown (near identical data was obtained for other replicates, data not shown).**

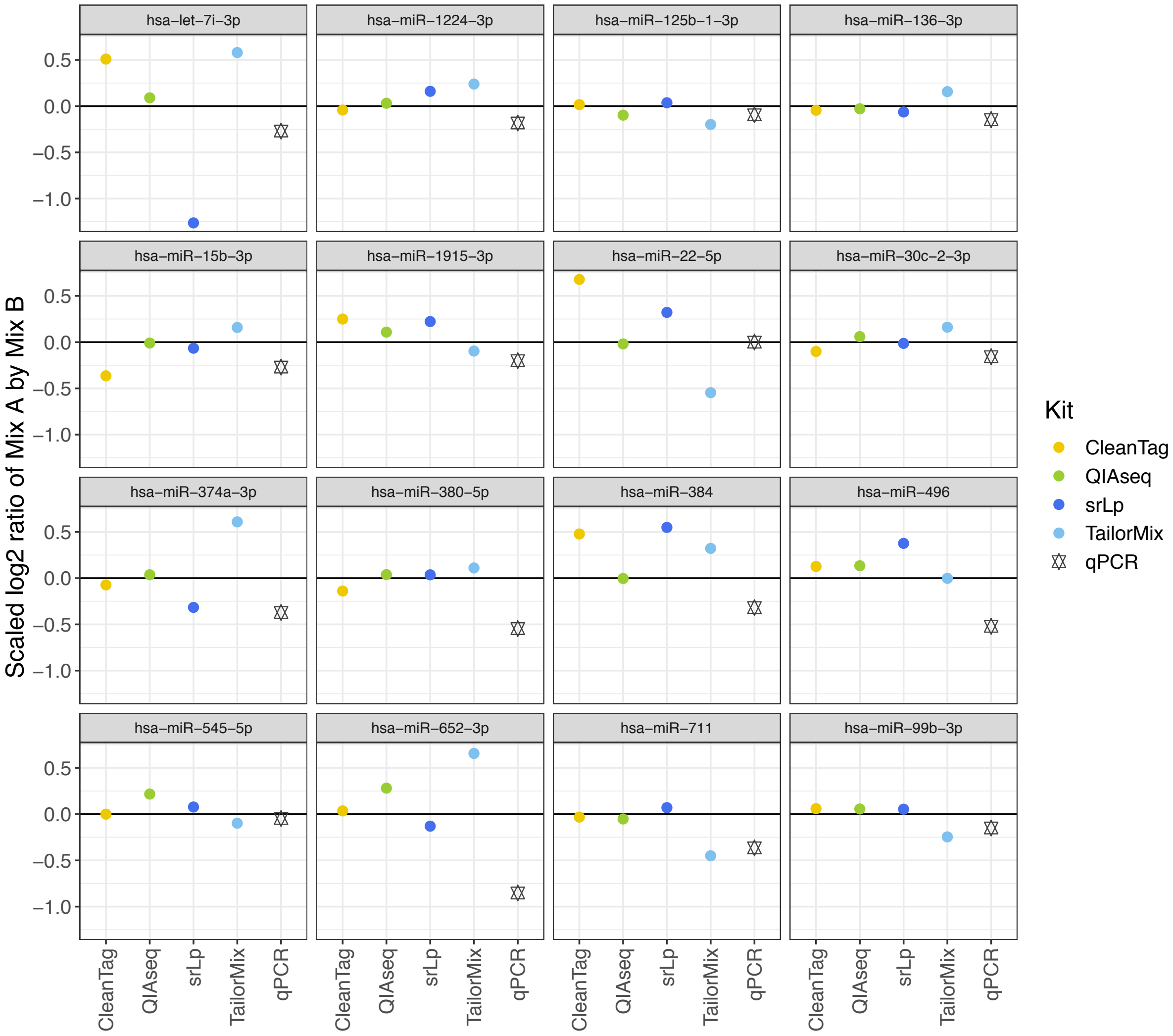


**Figure S14: Scaled and normalized log2 ratios of mix A and mix B for the read counts (sequencing data, represented by the library prep kits) and qPCR (copies detected relative to a standard curve) for 16 selected miRNAs. The horizontal black line represents the expected ratio.**

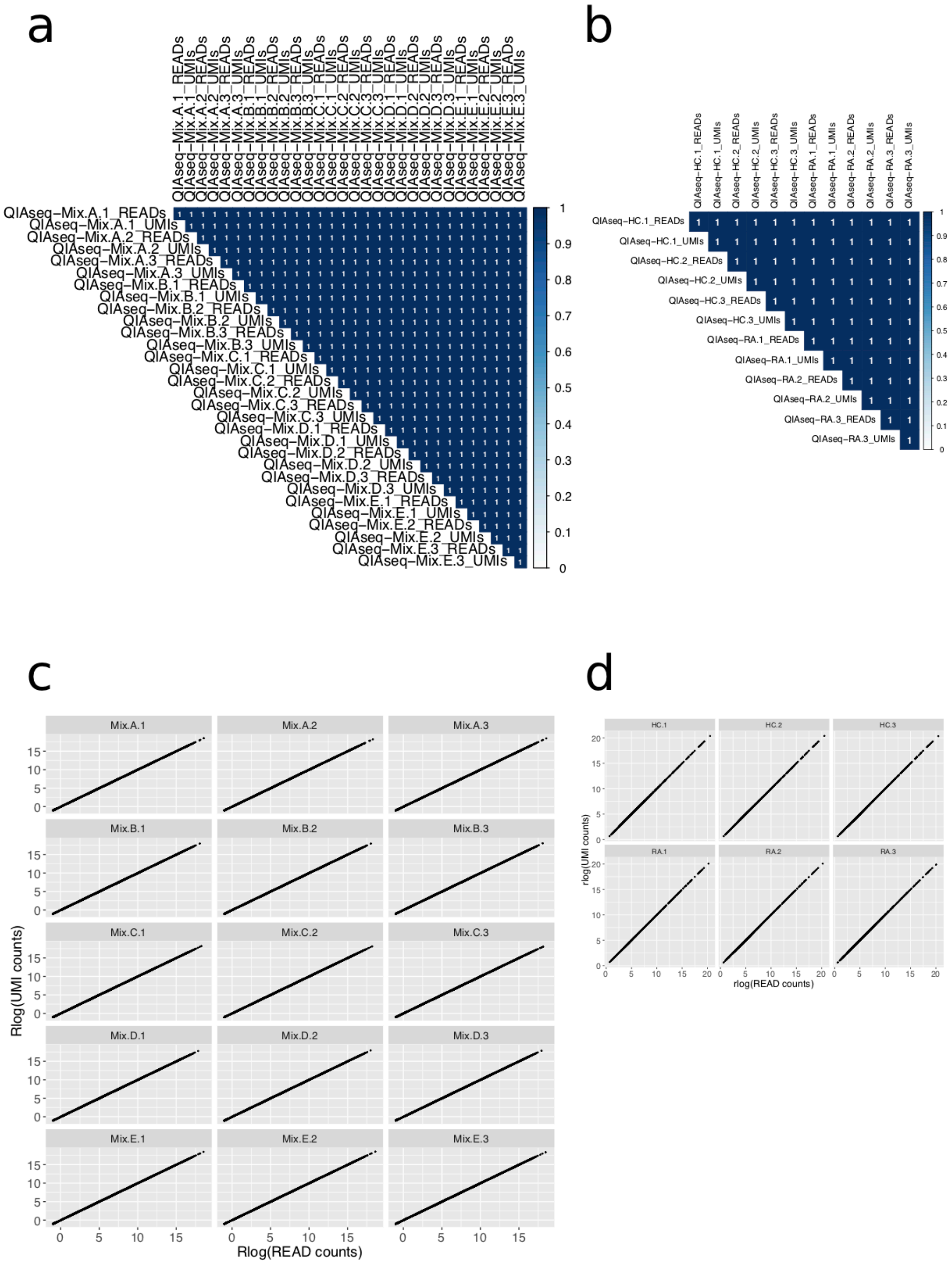


**Figure S15: Pearson correlation calculations and correlation plots between the rlog transformed UMI counts and the ordinary read counts for the synthetic miRNA samples (A, C) and the rheumatoid arthritis (RA) and healthy control (HC) human total RNA samples (B, D) of the QIAseq samples. The analysis tool GeneGlobe was used to create the UMI and read counts.**

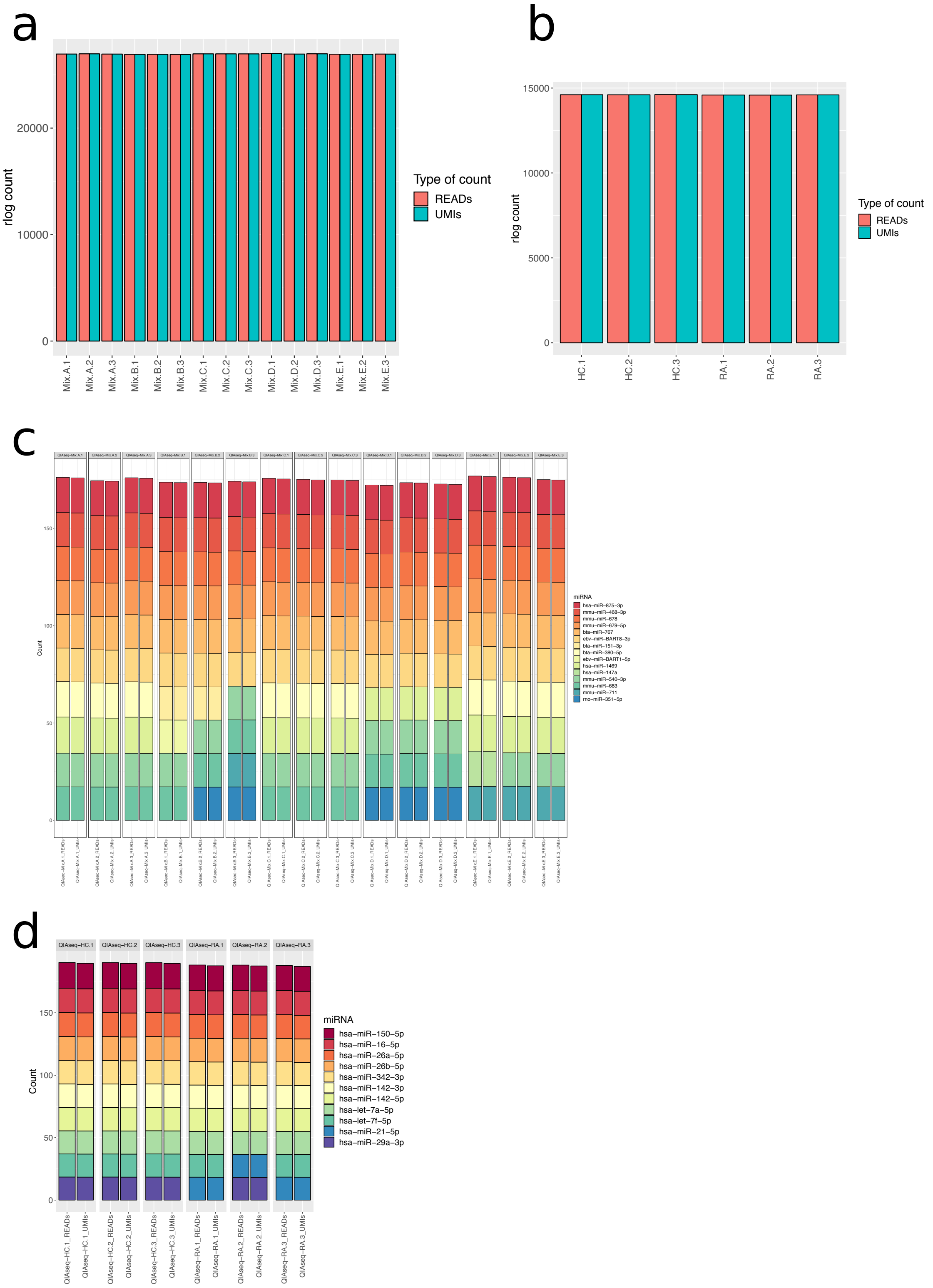


**Figure S16: QIAseq UMI analysis. Bar charts presenting the rlog sum of all UMI and ordinary reads counts in A: synthetic miRNA samples and B: human total RNA samples. Stacked bar plots present the ten most abundant ordinary rlog read counts and the UMI rlog count equivalent for C: synthetic miRNA samples and D: human total RNA samples.**
