## Supplementary Tables for "Systematic assessment of commercially available low-input miRNA library preparation kits"

S1 Table

### Table 1: Sequences and concentrations of the 40 non-equimolar miRNAs in the mixes A - E

| | Concentration [pmol] | | | | | |
|---|---|---|---|---|---|---|
| miRNA | MixA | MixB | MixC | MixD | MixE | Sequence |
| hsa-miR-380-5p | 300 | 3 | 225.75 | 77.25 | 30.00 | UGGUUGACCAUAGAACAUGCGC |
| hsa-miR-1224-3p | 300 | 3 | 225.75 | 77.25 | 30.00 | CCCCACCUCCUCUCUCCUCAG |
| hsa-miR-136-3p | 300 | 3 | 225.75 | 77.25 | 30.00 | CAUCAUCGUCUCAAAUGAGUCU |
| hsa-miR-1277-3p | 300 | 3 | 225.75 | 77.25 | 30.00 | UACGUAGAUAUAUAUGUAUUUU |
| hsa-miR-1469 | 300 | 3 | 225.75 | 77.25 | 30.00 | CUCGGCGCGGGGCGCGGGCUCC |
| hsa-miR-30c-2-3p | 120 | 12 | 93 | 39 | 12.00 | CUGGGAGAAGGCUGUUUACUCU |
| hsa-miR-139-3p | 120 | 12 | 93 | 39 | 12.00 | UGGAGACGCGGCCCUGUUGGAGU |
| hsa-miR-29a-5p | 120 | 12 | 93 | 39 | 12.00 | ACUGAUUUCUUUUGGUGUUCAG |
| hsa-miR-2052 | 120 | 12 | 93 | 39 | 12.00 | UGUUUUGAUAACAGUAAUGU |
| hsa-miR-1181 | 120 | 12 | 93 | 39 | 12.00 | CCGUCGCCGCCACCCGAGCCG |
| hsa-miR-652-3p | 60 | 12 | 48 | 24 | 6.00 | AAUGGCGCCACUAGGGUUGUG |
| hsa-miR-1307-3p | 60 | 12 | 48 | 24 | 6.00 | ACUCGGCGUGGCGUCGGUCGUG |
| hsa-miR-15b-3p | 60 | 12 | 48 | 24 | 6.00 | CGAAUCAUUAUUUGCUGCUCUA |
| hsa-miR-1252-5p | 60 | 12 | 48 | 24 | 6.00 | AGAAGGAAAUUGAAUUCAUUUA |
| hsa-miR-762 | 60 | 12 | 48 | 24 | 6.00 | GGGGCUGGGGCCGGGGCCGAGC |
| hsa-let-7i-3p | 60 | 30 | 52.5 | 37.5 | 6.00 | CUGCGCAAGCUACUGCCUUGCU |
| hsa-miR-1224-5p | 60 | 30 | 52.5 | 37.5 | 6.00 | GUGAGGACUCGGGAGGUGG |
| hsa-miR-153-5p | 60 | 30 | 52.5 | 37.5 | 6.00 | UCAUUUUUGUGAUGUUGCAGCU |
| hsa-miR-374a-3p | 60 | 30 | 52.5 | 37.5 | 6.00 | CUUAUCAGAUUGUAUUGUAAUU |
| hsa-miR-1199-5p | 60 | 30 | 52.5 | 37.5 | 6.00 | CCUGAGCCCGGGCCGCGCAG |
| hsa-miR-125a-3p | 30 | 60 | 37.5 | 52.5 | 3.00 | ACAGGUGAGGUUCUUGGGAGCC |
| hsa-miR-106b-3p | 30 | 60 | 37.5 | 52.5 | 3.00 | CCGCACUGUGGGUACUUGCUGC |
| hsa-miR-22-5p | 30 | 60 | 37.5 | 52.5 | 3.00 | AGUUCUUCAGUGGCAAGCUUUA |
| hsa-miR-384 | 30 | 60 | 37.5 | 52.5 | 3.00 | AUUCCUAGAAAUUGUUCAUA |
| hsa-miR-940 | 30 | 60 | 37.5 | 52.5 | 3.00 | AAGGCAGGGCCCCCGCUCCCC |
| hsa-miR-218-2-3p | 12 | 60 | 24 | 48 | 1.20 | CAUGGUUCUGUCAAGCACCGCG |
| hsa-miR-99b-3p | 12 | 60 | 24 | 48 | 1.20 | CAAGCUCGUGUCUGUGGGUCCG |
| hsa-miR-496 | 12 | 60 | 24 | 48 | 1.20 | UGAGUAUUACAUGGCCAAUCUC |
| hsa-miR-545-5p | 12 | 60 | 24 | 48 | 1.20 | UCAGUAAAUGUUUAUUAGAUGA |
| hsa-miR-718 | 12 | 60 | 24 | 48 | 1.20 | CUUCCGCCCCGCCGGGCGUCG |
| hsa-miR-1226-3p | 12 | 120 | 39 | 93 | 1.20 | UCACCAGCCCUGUGUUCCCUAG |
| hsa-miR-326 | 12 | 120 | 39 | 93 | 1.20 | CCUCUGGGCCCUUCCUCCAG |
| hsa-miR-802 | 12 | 120 | 39 | 93 | 1.20 | CAGUAACAAAGAUUCAUCCUUGU |
| hsa-miR-190a-3p | 12 | 120 | 39 | 93 | 1.20 | CUAUAUAUCAAACAUAUUCCU |
| hsa-miR-1915-3p | 12 | 120 | 39 | 93 | 1.20 | CCCCAGGGCGACGCGGCGGG |
| hsa-miR-125b-1-3p | 3 | 300 | 77.25 | 225.75 | 0.30 | ACGGGUUAGGCUCUUGGGAGCU |
| hsa-miR-711 | 3 | 300 | 77.25 | 225.75 | 0.30 | GGGACCCAGGGAGAGACGUAAG |
| hsa-miR-1197 | 3 | 300 | 77.25 | 225.75 | 0.30 | UAGGACACAUGGUCUACUUCU |

| | | | | | | |
|---|---|---|---|---|---|---|
| hsa-miR-2054 | 3 | 300 | 77.25 | 225.75 | 0.30 | CUGUAAUAUAAAUUUAAUUUAUU |
| hsa-miR-1470 | 3 | 300 | 77.25 | 225.75 | 0.30 | GCCCUCCGCCCGUGCACCCCG |

### S2 Table

**Table 2: Summary of sample material.**

| Continous sample number | Sample Amount *(miRNA/yeast total RNA) **(Total human RNA) | Sample Type | Mix Type / state | Replicate | Index Sequence | Common (Illumina) index name |
|---|---|---|---|---|---|---|
| 1 | 1 ng / 9 ng* | Synthetic miRNA | A | 1 | ATCACG | TruSeq Index 1 |
| 2 | 1 ng / 9 ng* | Synthetic miRNA | A | 2 | CGATGT | TruSeq Index 2 |
| 3 | 1 ng / 9 ng* | Synthetic miRNA | A | 3 | TTAGGC | TruSeq Index 3 |
| 4 | 1 ng / 9 ng* | Synthetic miRNA | B | 1 | TGACCA | TruSeq Index 4 |
| 5 | 1 ng / 9 ng* | Synthetic miRNA | B | 2 | ACAGTG | TruSeq Index 5 |
| 6 | 1 ng / 9 ng* | Synthetic miRNA | B | 3 | GCCAAT | TruSeq Index 6 |
| 7 | 1 ng / 9 ng* | Synthetic miRNA | C | 1 | CAGATC | TruSeq Index 7 |
| 8 | 1 ng / 9 ng* | Synthetic miRNA | C | 2 | ACTTGA | TruSeq Index 8 |
| 9 | 1 ng / 9 ng* | Synthetic miRNA | C | 3 | GATCAG | TruSeq Index 9 |
| 10 | 1 ng / 9 ng* | Synthetic miRNA | D | 1 | TAGCTT | TruSeq Index 10 |
| 11 | 1 ng / 9 ng* | Synthetic miRNA | D | 2 | GGCTAC | TruSeq Index 11 |
| 12 | 1 ng / 9 ng* | Synthetic miRNA | D | 3 | CTTGTA | TruSeq Index 12 |
| 13 | 0.1 ng / 9.9 ng* | Synthetic miRNA | E | 1 | AGTCAA | TruSeq Index 13 |
| 14 | 0.1 ng / 9.9 ng* | Synthetic miRNA | E | 2 | AGTTCC | TruSeq Index 14 |
| 15 | 0.1 ng / 9.9 ng* | Synthetic miRNA | E | 3 | ATGTCA | TruSeq Index 15 |
| 16 | 100 ng** | Total human RNA | Disease state | 1 | CCGTCC | TruSeq Index 16 |
| 17 | 100 ng** | Total human RNA | Disease state | 2 | GTCCGC | TruSeq Index 18 |

| | | | | | | |
|---|---|---|---|---|---|---|
| 18 | 100 ng** | Total human RNA | Disease state | 3 | GTGAAA | TruSeq Index 19 |
| 19 | 100 ng** | Total human RNA | Control state | 1 | GTGGCC | TruSeq Index 20 |
| 20 | 100 ng** | Total human RNA | Control state | 2 | GTTTCG | TruSeq Index 21 |
| 21 | 100 ng** | Total human RNA | Control state | 3 | CGTACG | TruSeq Index 22 |

**S3 Table**

**Table 3: miRXplore Universal Reference miRNAs which had total sequence accordance with sequences in the *Saccharomyces cerevisae* (sacCer3) genome**

| miRNA | Sequence |
|---|---|
| EBV-12 | UCCUGUGGUGUUUGGUGUGGUU |
| EBV-15 | GUCAGUGGUUUUGUUUCCUUGA |
| EBV-19-3P | UUUUGUUUGCUUGGGAAUGCU |
| LET-7I | UGAGGUAGUAGUUUGUGCUGUU |
| MIR-1-rno | UGGAAUGUAAAGAAGUGUGUAU |
| MIR-135A | UAUGGCUUUUUAUUCCUAUGUGA |
| MIR-142-3P | UGUAGUGUUUCCUACUUUAUGGA |
| MIR-144 | UACAGUAUAGAUGAUGUACU |
| MIR-190 | UGAUAUGUUUGAUAUAUUAGGU |
| MIR-190B | UGAUAUGUUUGAUAUUGGGUU |
| MIR-190B-rno | UGAUAUGUUUGAUAUUAGGUU |
| MIR-202-5P | UUUCCUAUGCAUAUACUUCUUU |
| MIR-204 | UUCCCUUUGUCAUCCUAUGCCU |
| MIR-211 | UUCCCUUUGUCAUCCUUUGCCU |
| MIR-291B-3P-mmu | AAAGUGCAUCCAUUUUGUUUGU |
| MIR-291B-3P-rno | AAAGUGCAUCCAUUUUGUUAGU |
| MIR-297A | AUGUAUGUGUGCAUGUGCAUGU |
| MIR-322 | CAGCAGCAAUUCAUGUUUUGGA |
| MIR-322-hsa | CAGCAGCAAUUCAUGUUUUGAA |
| MIR-333 | GUGGUGUGCUAGUUACUUUU |
| MIR-337-3P-hsa | CUCCUAUAUGAUGCCUUUCUUC |
| MIR-33A | GUGCAUUGUAGUUGCAUUGCA |
| MIR-33B | GUGCAUUGCUGUUGCAUUGC |
| MIR-369-3P | AAUAAUACAUGGUUGAUCUUU |
| MIR-374-star | GGUUGUAUUAUCAUUGUCCGAG |
| MIR-376B-star | GUGGAUAUUCCUUCUAUGGUU |
| MIR-380-3P-hsa | UAUGUAAUAUGGUCCACAUCUU |
| MIR-384-3P-hsa | AUUCCUAGAAAUUGUUCAUAAU |
| MIR-421-5P | GGCCUCAUUAAAUGUUUGUUG |
| MIR-450A-5P | UUUUGCGAUGUGUUCCUAAUAU |
| MIR-450B-5P-hsa | UUUUGCAAUAUGUUCCUGAAUA |
| MIR-450B-5P-mmu | UUUUGCAGUAUGUUCCUGAAUA |
| MIR-450B-5P-rno | UUUUGCAGUAUGUUCCUGUAUA |
| MIR-466C-5P-rno | UGUGAUGUGUGCAUGUACAUG |

| | |
|---|---|
| MIR-467E | AUAAGUGUGAGCAUGUAUAUGU |
| MIR-488-hsa | UUGAAAGGCUAUUUCUUGGUCU |
| MIR-507 | UUUUGCACCUUUUGGAGUGAA |
| MIR-509-3P-mmu | UGAUUGACAUUUCUGUAAUGG |
| MIR-516A-3P | UGCUUCCUUUCAGAGGGU |
| MIR-547-mmu | CUUGGUACAUCUUUGAGUGAGU |
| MIR-548C-5P | AAAAGUAAUUGCGGUUUUUGCC |
| MIR-553 | AAAACGGUGAGAUUUUGUUUU |
| MIR-581 | UCUUGUGUUCUCUAGAUCAGU |
| MIR-586 | UAUGCAUUGUAUUUUUAGGUCC |
| MIR-590-3P | UAAUUUUAUGUAUAAGCUAGU |
| MIR-590-3P-rno | UAAUUUUAUGUAUGAGCUGGU |
| MIR-603 | CACACACUGCAAUUACUUUUGC |
| MIR-607 | GUUCAAAUCCAGAUCUAUAAC |
| MIR-609 | AGGGUGUUUCUCUCAUCUCU |
| MIR-620 | AUGGAGAUAGAUAUAGAAAU |
| MIR-651 | UUUAGGAUAAGCUUGACUUUUG |
| MIR-669B | AGUUUUGUGUGCAUGUGCAUGU |
| MIR-670 | AUCCCUGAGUGUAUGUGGUGAA |
| MIR-684 | AGUUUUCCCUUCAAGUCAA |
| MIR-698 | CAUUCUCGUUUCCUUCCCU |
| MIR-7 | UGGAAGACUAGUGAUUUUGUUGU |
| MIR-876-5P-mmu | UGGAUUUCUCUGUGAAUCACUA |
| MIR-881-mmu | AACUGUGUCUUUUCUGAAUAGA |
| MIR-98 | UGAGGUAGUAAGUUGUAUUGUU |

S4 Table

Table 4: Overview of sample names, replicates, mix and the number of reads for the different bioinformatics steps

| Participant name | Kit_name | Kit name used in this study | Replicate | Mix | HiSeq Lane | Raw read count | # reads after trimming | # reads after mapping | # reads after counting = miRNA reads |
|---|---|---|---|---|---|---|---|---|---|
| Diagenode-1 | CATS Small RNA-Seq Kit | CATS | 1 | A | L001 | 1044990 | 983726 | 464342 | 210874 |
| Diagenode-2 | CATS Small RNA-Seq Kit | CATS | 2 | A | L002 | 1657065 | 1557685 | 775521 | 350790 |
| Diagenode-3 | CATS Small RNA-Seq Kit | CATS | 3 | A | L003 | 1040559 | 980308 | 454236 | 204194 |
| Diagenode-4 | CATS Small RNA-Seq Kit | CATS | 1 | B | L004 | 1012071 | 963230 | 492407 | 223976 |
| Diagenode-5 | CATS Small RNA-Seq Kit | CATS | 2 | B | L005 | 1368175 | 1314283 | 594308 | 234985 |
| Diagenode-6 | CATS Small RNA-Seq Kit | CATS | 3 | B | L001 | 1792546 | 1721449 | 744133 | 272768 |
| Diagenode-7 | CATS Small RNA-Seq Kit | CATS | 1 | C | L002 | 1427147 | 1363859 | 681475 | 253041 |
| Diagenode-8 | CATS Small RNA-Seq Kit | CATS | 2 | C | L003 | 1521215 | 1444048 | 740731 | 266561 |
| Diagenode-9 | CATS Small RNA-Seq Kit | CATS | 3 | C | L004 | 1220417 | 1138980 | 545140 | 185295 |
| Diagenode-10 | CATS Small RNA-Seq Kit | CATS | 1 | D | L005 | 4522878 | 4226522 | 2175091 | 775901 |
| Diagenode-11 | CATS Small RNA-Seq Kit | CATS | 2 | D | L001 | 6184503 | 5821807 | 3319459 | 1214419 |
| Diagenode-12 | CATS Small RNA-Seq Kit | CATS | 3 | D | L002 | 4520202 | 4260836 | 2368897 | 878785 |
| Diagenode-13 | CATS Small RNA-Seq Kit | CATS | 1 | E | L003 | 4053616 | 3501644 | 791565 | 283123 |
| Diagenode-14 | CATS Small RNA-Seq Kit | CATS | 2 | E | L004 | 3319238 | 2934993 | 805353 | 293457 |
| Diagenode-15 | CATS Small RNA-Seq Kit | CATS | 3 | E | L005 | 4047587 | 3442401 | 537837 | 177740 |
| Diagenode-16 | CATS Small RNA-Seq Kit | CATS | 1 | RA | L001 | 5118079 | 3655276 | 55660 | 11641 |
| Diagenode-17 | CATS Small RNA-Seq Kit | CATS | 2 | RA | L002 | 4421600 | 2984739 | 50605 | 9556 |
| Diagenode-18 | CATS Small RNA-Seq Kit | CATS | 3 | RA | L003 | 4316233 | 3055548 | 49831 | 10198 |
| Diagenode-19 | CATS Small RNA-Seq Kit | CATS | 1 | HC | L004 | 3317101 | 2449536 | 35597 | 8139 |
| Diagenode-20 | CATS Small RNA-Seq Kit | CATS | 2 | HC | L005 | 3844218 | 2795815 | 43328 | 9667 |
| Diagenode-21 | CATS Small RNA-Seq Kit | CATS | 3 | HC | L001 | 4460329 | 3174695 | 48767 | 10671 |

| | | | | | | | | | |
|---|---|---|---|---|---|---|---|---|---|
| Lexogen-1 | Small RNA-Seq Library Prep Kit | srLp | 1 | A | L005 | 19970677 | 19824293 | 15392907 | 14559791 |
| Lexogen-2 | Small RNA-Seq Library Prep Kit | srLp | 2 | A | L001 | 20295024 | 20124545 | 15820320 | 14971660 |
| Lexogen-3 | Small RNA-Seq Library Prep Kit | srLp | 3 | A | L002 | 24152499 | 23964309 | 18737467 | 17742053 |
| Lexogen-4 | Small RNA-Seq Library Prep Kit | srLp | 1 | B | L003 | 24648583 | 24482843 | 19106583 | 18117915 |
| Lexogen-5 | Small RNA-Seq Library Prep Kit | srLp | 2 | B | L004 | 14348353 | 14122166 | 10859771 | 10221601 |
| Lexogen-6 | Small RNA-Seq Library Prep Kit | srLp | 3 | B | L005 | 25621732 | 25439113 | 19574381 | 18516687 |
| Lexogen-7 | Small RNA-Seq Library Prep Kit | srLp | 1 | C | L001 | 25767806 | 25570256 | 20056292 | 18974321 |
| Lexogen-8 | Small RNA-Seq Library Prep Kit | srLp | 2 | C | L002 | 24589126 | 24417839 | 19137165 | 18113004 |
| Lexogen-9 | Small RNA-Seq Library Prep Kit | srLp | 3 | C | L003 | 25762284 | 25597563 | 20003077 | 18933822 |
| Lexogen-10 | Small RNA-Seq Library Prep Kit | srLp | 1 | D | L004 | 20524417 | 20351673 | 15591334 | 14736928 |
| Lexogen-11 | Small RNA-Seq Library Prep Kit | srLp | 2 | D | L005 | 22560264 | 22415200 | 17353731 | 16408453 |
| Lexogen-12 | Small RNA-Seq Library Prep Kit | srLp | 3 | D | L001 | 21708163 | 21525355 | 16953966 | 16014003 |
| Lexogen-13 | Small RNA-Seq Library Prep Kit | srLp | 1 | E | L002 | 21261063 | 20435057 | 13690962 | 12936140 |
| Lexogen-14 | Small RNA-Seq Library Prep Kit | srLp | 2 | E | L003 | 21349913 | 20761962 | 14130952 | 13297815 |
| Lexogen-15 | Small RNA-Seq Library Prep Kit | srLp | 3 | E | L004 | 20030874 | 19299274 | 12526116 | 11830778 |

| | | | | | | | | | |
|---|---|---|---|---|---|---|---|---|---|
| Lexogen-16 | Small RNA-Seq Library Prep Kit | srLp | 1 | RA | L005 | 4637425 | 4447693 | 1221499 | 758388 |
| Lexogen-17 | Small RNA-Seq Library Prep Kit | srLp | 2 | RA | L001 | 11145100 | 10706550 | 3211892 | 1983800 |
| Lexogen-18 | Small RNA-Seq Library Prep Kit | srLp | 3 | RA | L002 | 5994469 | 5505003 | 1413515 | 863268 |
| Lexogen-19 | Small RNA-Seq Library Prep Kit | srLp | 1 | HC | L003 | 10771427 | 9979458 | 2643810 | 1638276 |
| Lexogen-20 | Small RNA-Seq Library Prep Kit | srLp | 2 | HC | L004 | 13006115 | 11776620 | 3036805 | 1864069 |
| Lexogen-21 | Small RNA-Seq Library Prep Kit | srLp | 3 | HC | L005 | 8334900 | 7414996 | 2114300 | 1284163 |
| Qiagen-1 | QIAseq miRNA Library | QIAseq | 1 | A | L004 | 23185265 | 23105721 | 18777291 | 17696215 |
| Qiagen-2 | QIAseq miRNA Library | QIAseq | 2 | A | L005 | 26756917 | 26647535 | 21433321 | 20171612 |
| Qiagen-3 | QIAseq miRNA Library | QIAseq | 3 | A | L001 | 28887780 | 28792146 | 23309760 | 21996842 |
| Qiagen-4 | QIAseq miRNA Library | QIAseq | 1 | B | L002 | 23085124 | 23017717 | 18578204 | 17592394 |
| Qiagen-5 | QIAseq miRNA Library | QIAseq | 2 | B | L003 | 25025406 | 24942149 | 19993601 | 18929903 |
| Qiagen-6 | QIAseq miRNA Library | QIAseq | 3 | B | L004 | 29561642 | 29475996 | 23664866 | 22443650 |
| Qiagen-7 | QIAseq miRNA Library | QIAseq | 1 | C | L005 | 31710211 | 31605868 | 25433312 | 24014129 |
| Qiagen-8 | QIAseq miRNA Library | QIAseq | 2 | C | L001 | 30480862 | 30383218 | 24632401 | 23265808 |
| Qiagen-9 | QIAseq miRNA Library | QIAseq | 3 | C | L002 | 29744870 | 29640598 | 23979920 | 22630500 |
| Qiagen-10 | QIAseq miRNA Library | QIAseq | 1 | D | L003 | 33705683 | 33594072 | 27046929 | 25573975 |
| Qiagen-11 | QIAseq miRNA Library | QIAseq | 2 | D | L004 | 20481668 | 20429994 | 16463354 | 15592672 |
| Qiagen-12 | QIAseq miRNA Library | QIAseq | 3 | D | L005 | 24773931 | 24702652 | 19766974 | 18700148 |
| Qiagen-13 | QIAseq miRNA Library | QIAseq | 1 | E | L001 | 19217805 | 19145242 | 15291685 | 14446124 |
| Qiagen-14 | QIAseq miRNA Library | QIAseq | 2 | E | L002 | 19584832 | 19502838 | 15588051 | 14770461 |
| Qiagen-15 | QIAseq miRNA Library | QIAseq | 3 | E | L003 | 19004068 | 18939751 | 15127089 | 14340907 |
| Qiagen-16 | QIAseq miRNA Library | QIAseq | 1 | RA | L004 | 16750249 | 15365730 | 6331862 | 3622355 |
| Qiagen-17 | QIAseq miRNA Library | QIAseq | 2 | RA | L005 | 16516348 | 15147604 | 6289024 | 3585493 |

| | | | | | | | | | |
|---|---|---|---|---|---|---|---|---|---|
| Qiagen-18 | QIAseq miRNA Library | QIAseq | 3 | RA | L001 | 19347339 | 17694009 | 7402777 | 4154028 |
| Qiagen-19 | QIAseq miRNA Library | QIAseq | 1 | HC | L002 | 17571917 | 15767429 | 6474229 | 3677697 |
| Qiagen-20 | QIAseq miRNA Library | QIAseq | 2 | HC | L003 | 19037948 | 16131519 | 6682764 | 3796842 |
| Qiagen-21 | QIAseq miRNA Library | QIAseq | 3 | HC | L004 | 15540227 | 13750211 | 5645066 | 3171607 |
| SeqMatic-1 | TailorMix miRNA Sample Preparation Kit V3 | TailorMix | 1 | A | L003 | 15684848 | 15677220 | 12890832 | 12433820 |
| SeqMatic-2 | TailorMix miRNA Sample Preparation Kit V3 | TailorMix | 2 | A | L004 | 11749782 | 11739361 | 9607236 | 9255083 |
| SeqMatic-3 | TailorMix miRNA Sample Preparation Kit V3 | TailorMix | 3 | A | L005 | 11781326 | 11774607 | 9617259 | 9270709 |
| SeqMatic-4 | TailorMix miRNA Sample Preparation Kit V3 | TailorMix | 1 | B | L001 | 11477845 | 11458927 | 9286460 | 8970365 |
| SeqMatic-5 | TailorMix miRNA Sample Preparation Kit V3 | TailorMix | 2 | B | L002 | 12561555 | 12550122 | 10180334 | 9798864 |
| SeqMatic-6 | TailorMix miRNA Sample Preparation Kit V3 | TailorMix | 3 | B | L003 | 11473824 | 11466086 | 9254400 | 8919391 |
| SeqMatic-7 | TailorMix miRNA Sample Preparation Kit V3 | TailorMix | 1 | C | L004 | 9880077 | 9872311 | 8094232 | 7788482 |
| SeqMatic-8 | TailorMix miRNA Sample Preparation Kit V3 | TailorMix | 2 | C | L005 | 10731731 | 10718419 | 8741884 | 8233840 |
| SeqMatic-9 | TailorMix miRNA Sample Preparation Kit V3 | TailorMix | 3 | C | L001 | 12921035 | 12912821 | 10540960 | 10155065 |
| SeqMatic-10 | TailorMix miRNA Sample Preparation Kit V3 | TailorMix | 1 | D | L002 | 14442889 | 14436274 | 11655182 | 11216094 |
| SeqMatic-11 | TailorMix miRNA Sample Preparation Kit V3 | TailorMix | 2 | D | L003 | 12948439 | 12941050 | 10569444 | 10165416 |
| SeqMatic-12 | TailorMix miRNA Sample Preparation Kit V3 | TailorMix | 3 | D | L004 | 12904412 | 12894631 | 10540604 | 10139354 |
| SeqMatic-13 | TailorMix miRNA Sample Preparation Kit V3 | TailorMix | 1 | E | L005 | 12950192 | 12871478 | 10480434 | 10055674 |

| | | | | | | | | | |
|---|---|---|---|---|---|---|---|---|---|
| SeqMatic-14 | TailorMix miRNA Sample Preparation Kit V3 | TailorMix | 2 | E | L001 | 16109251 | 16072149 | 13294638 | 12777666 |
| SeqMatic-15 | TailorMix miRNA Sample Preparation Kit V3 | TailorMix | 3 | E | L002 | 18158394 | 18066346 | 14834083 | 14242970 |
| SeqMatic-16 | TailorMix miRNA Sample Preparation Kit V3 | TailorMix | 1 | RA | L003 | 11401076 | 11197308 | 4821171 | 3162024 |
| SeqMatic-17 | TailorMix miRNA Sample Preparation Kit V3 | TailorMix | 2 | RA | L004 | 12350057 | 12073599 | 5875470 | 3404519 |
| SeqMatic-18 | TailorMix miRNA Sample Preparation Kit V3 | TailorMix | 3 | RA | L005 | 14445783 | 14100175 | 7060962 | 4557694 |
| SeqMatic-19 | TailorMix miRNA Sample Preparation Kit V3 | TailorMix | 1 | HC | L001 | 11222613 | 11011398 | 5014127 | 3237729 |
| SeqMatic-20 | TailorMix miRNA Sample Preparation Kit V3 | TailorMix | 2 | HC | L002 | 11216132 | 11024639 | 5418281 | 3457010 |
| SeqMatic-21 | TailorMix miRNA Sample Preparation Kit V3 | TailorMix | 3 | HC | L003 | 12926192 | 12729343 | 5522928 | 3628822 |
| Takara-1 | SMARTer® miRNA-seq Kit (Beta version) | SMARTer | 1 | A | L006 | 7023334 | 6962533 | 195609 | 0 |
| Takara-2 | SMARTer® miRNA-seq Kit (Beta version) | SMARTer | 2 | A | L006 | 6753690 | 6669061 | 276826 | 4 |
| Takara-3 | SMARTer® miRNA-seq Kit (Beta version) | SMARTer | 3 | A | L006 | 4817693 | 4768804 | 149947 | 2 |
| Takara-4 | SMARTer® miRNA-seq Kit (Beta version) | SMARTer | 1 | B | L006 | 6467640 | 6377764 | 261651 | 1 |
| Takara-5 | SMARTer® miRNA-seq Kit (Beta version) | SMARTer | 2 | B | L006 | 4371221 | 4312673 | 119359 | 2 |
| Takara-6 | SMARTer® miRNA-seq Kit (Beta version) | SMARTer | 3 | B | L006 | 1542657 | 1522402 | 16596 | 0 |
| Takara-7 | SMARTer® miRNA-seq Kit (Beta version) | SMARTer | 1 | C | L006 | 4403705 | 4326034 | 154247 | 0 |

| | | | | | | | | | |
|---|---|---|---|---|---|---|---|---|---|
| Takara-8 | SMARTer® miRNA-seq Kit (Beta version) | SMARTer | 2 | C | L006 | 2821746 | 2778865 | 74044 | 1 |
| Takara-9 | SMARTer® miRNA-seq Kit (Beta version) | SMARTer | 3 | C | L006 | 2102677 | 2067983 | 57359 | 1 |
| Takara-10 | SMARTer® miRNA-seq Kit (Beta version) | SMARTer | 1 | D | L006 | 1168082 | 1149648 | 24573 | 0 |
| Takara-11 | SMARTer® miRNA-seq Kit (Beta version) | SMARTer | 2 | D | L006 | 7350966 | 7304109 | 297016 | 4 |
| Takara-12 | SMARTer® miRNA-seq Kit (Beta version) | SMARTer | 3 | D | L006 | 5835162 | 5801837 | 281379 | 1 |
| Takara-13 | SMARTer® miRNA-seq Kit (Beta version) | SMARTer | 1 | E | L006 | 5091337 | 5058843 | 252277 | 7 |
| Takara-14 | SMARTer® miRNA-seq Kit (Beta version) | SMARTer | 2 | E | L006 | 8755804 | 8702278 | 403984 | 4 |
| Takara-15 | SMARTer® miRNA-seq Kit (Beta version) | SMARTer | 3 | E | L006 | 3964212 | 3930005 | 198341 | 4 |
| Takara-16 | SMARTer® miRNA-seq Kit (Beta version) | SMARTer | 1 | RA | L006 | 1272993 | 1184422 | 23839 | 2 |
| Takara-17 | SMARTer® miRNA-seq Kit (Beta version) | SMARTer | 2 | RA | L006 | 736326 | 693272 | 8460 | 3 |
| Takara-18 | SMARTer® miRNA-seq Kit (Beta version) | SMARTer | 3 | RA | L006 | 812827 | 764140 | 12442 | 1 |
| Takara-19 | SMARTer® miRNA-seq Kit (Beta version) | SMARTer | 1 | HC | L006 | 424237 | 408105 | 5865 | 0 |
| Takara-20 | SMARTer® miRNA-seq Kit (Beta version) | SMARTer | 2 | HC | L006 | 519032 | 500373 | 5840 | 1 |
| Takara-21 | SMARTer® miRNA-seq Kit (Beta version) | SMARTer | 3 | HC | L006 | 693605 | 667391 | 9598 | 4 |
| TriLink-1 | CleanTag™ Small RNA Library Prep Kit | CleanTag | 1 | A | L002 | 7593966 | 7564341 | 6053589 | 5600246 |

| | | | | | | | | | |
|---|---|---|---|---|---|---|---|---|---|
| TriLink-2 | CleanTag™ Small RNA Library Prep Kit | CleanTag | 2 | A | L004 | 9142144 | 9105473 | 7083966 | 6552021 |
| TriLink-3 | CleanTag™ Small RNA Library Prep Kit | CleanTag | 3 | A | L005 | 8749658 | 8718113 | 6724519 | 6215798 |
| TriLink-4 | CleanTag™ Small RNA Library Prep Kit | CleanTag | 1 | B | L001 | 10681905 | 10639525 | 8319339 | 7722102 |
| TriLink-5 | CleanTag™ Small RNA Library Prep Kit | CleanTag | 2 | B | L002 | 4197732 | 4104810 | 3109754 | 2789056 |
| TriLink-6 | CleanTag™ Small RNA Library Prep Kit | CleanTag | 3 | B | L003 | 10154654 | 10116255 | 7955756 | 7418087 |
| TriLink-7 | CleanTag™ Small RNA Library Prep Kit | CleanTag | 1 | C | L004 | 11545893 | 11503978 | 9138364 | 8499074 |
| TriLink-8 | CleanTag™ Small RNA Library Prep Kit | CleanTag | 2 | C | L005 | 9624896 | 9573677 | 7642335 | 7092470 |
| TriLink-9 | CleanTag™ Small RNA Library Prep Kit | CleanTag | 3 | C | L001 | 10044117 | 10010797 | 7919993 | 7384238 |
| TriLink-10 | CleanTag™ Small RNA Library Prep Kit | CleanTag | 1 | D | L002 | 12220416 | 12161333 | 9677497 | 8995828 |
| TriLink-11 | CleanTag™ Small RNA Library Prep Kit | CleanTag | 2 | D | L003 | 8740509 | 8695883 | 6908065 | 6407309 |
| TriLink-12 | CleanTag™ Small RNA Library Prep Kit | CleanTag | 3 | D | L004 | 10724883 | 10680880 | 8519096 | 7937673 |
| TriLink-13 | CleanTag™ Small RNA Library Prep Kit | CleanTag | 1 | E | L005 | 9633137 | 9547013 | 7453570 | 6943194 |
| TriLink-14 | CleanTag™ Small RNA Library Prep Kit | CleanTag | 2 | E | L001 | 12705446 | 12571078 | 9670377 | 8958714 |
| TriLink-15 | CleanTag™ Small RNA Library Prep Kit | CleanTag | 3 | E | L002 | 10822925 | 10717407 | 8450515 | 7881159 |
| TriLink-16 | CleanTag™ Small RNA Library Prep Kit | CleanTag | 1 | RA | L003 | 16246499 | 15030847 | 2252856 | 1463296 |

| | | | | | | | | | |
|---|---|---|---|---|---|---|---|---|---|
| TriLink-17 | CleanTag™ Small RNA Library Prep Kit | CleanTag | 2 | RA | L004 | 16464052 | 15462438 | 2031962 | 1320081 |
| TriLink-18 | CleanTag™ Small RNA Library Prep Kit | CleanTag | 3 | RA | L005 | 19822754 | 18607039 | 2503500 | 1633274 |
| TriLink-19 | CleanTag™ Small RNA Library Prep Kit | CleanTag | 1 | HC | L003 | 19473071 | 18602100 | 1745644 | 1161503 |
| TriLink-20 | CleanTag™ Small RNA Library Prep Kit | CleanTag | 2 | HC | L001 | 28161853 | 26494954 | 2785144 | 1817911 |
| TriLink-21 | CleanTag™ Small RNA Library Prep Kit | CleanTag | 3 | HC | L002 | 25667555 | 24262971 | 2771924 | 1819800 |

S5 Table

Table 5: Intra-rater reliability of the three library preparation kit replicates of Mix A to Mix E and , RA and healthy control samples measured by ICC(3,1) and absolute agreement

| Sample | ICC values | srLp | QIAseq | TailorMix | CleanTaq |
|---|---|---|---|---|---|
| | ICC | **0.995** | **0.996** | **0.995** | **0.996** |
| Mix A | lbound | 0.994 | 0.996 | 0.994 | 0.995 |
| | ubound | 0.995 | 0.997 | 0.995 | 0.996 |
| | ICC | **0.995** | **0.997** | **0.992** | **0.992** |
| Mix B | lbound | 0.994 | 0.997 | 0.992 | 0.991 |
| | ubound | 0.995 | 0.997 | 0.993 | 0.993 |
| | ICC | **0.995** | **0.997** | **0.984** | **0.995** |
| Mix C | lbound | 0.995 | 0.997 | 0.982 | 0.994 |
| | ubound | 0.996 | 0.998 | 0.985 | 0.995 |
| | ICC | **0.994** | **0.996** | **0.996** | **0.991** |
| Mix D | lbound | 0.994 | 0.996 | 0.996 | 0.990 |
| | ubound | 0.995 | 0.997 | 0.997 | 0.992 |
| | ICC | **0.992** | **0.994** | **0.994** | **0.988** |
| Mix E | lbound | 0.991 | 0.994 | 0.994 | 0.987 |
| | ubound | 0.993 | 0.995 | 0.995 | 0,990 |
| Rheumatoid | ICC | **0.990** | **0.992** | **0.991** | **0.994** |
| arthritis | lbound | 0.989 | 0.991 | 0.990 | 0.993 |
| patients | ubound | 0.991 | 0.992 | 0.992 | 0.995 |
| | ICC | **0.990** | **0.994** | **0.993** | **0.994** |
| Healthy | lbound | 0.989 | 0.994 | 0.992 | 0.994 |
| controls | ubound | 0.991 | 0.995 | 0.993 | 0.995 |

lbound = lower confidence interval bound; ubound = upper confidence interval bound

S6 Table:

Table 6: Inter-rater reliability of the first replicate of each library preparation kit in Mix A to Mix E, RA and healthy control samples measured by ICC(3,1) and absolute agreement.

| Sample | ICC | lBound | uBound |
|---|---|---|---|
| Mix A | **0.841** | 0.826 | 0.855 |
| Mix B | **0.837** | 0.822 | 0.852 |
| Mix C | **0.837** | 0.821 | 0.851 |
| Mix D | **0.83** | 0.814 | 0.845 |
| Mix E | **0.839** | 0.824 | 0.854 |
| Rheumatoid arthritis patient | **0.964** | 0.961 | 0.967 |
| Healthy controls | **0.955** | 0.951 | 0.959 |

lbound = lower confidence interval bound; ubound = upper confidence interval bound

S7 Table

Table 7: Within kit coefficient of variation of the 903 equimolar miRNA oligonucleotides

| Mix | Kit | Replicate | Coefficient of variation |
|---|---|---|---|
| A | srlP | 1 | 40.85 |
| A | srlP | 2 | 41.04 |
| A | srlP | 3 | 40.99 |
| B | srlP | 1 | 40.88 |
| B | srlP | 2 | 40.75 |
| B | srlP | 3 | 40.95 |
| C | srlP | 1 | 41.00 |
| C | srlP | 2 | 41.05 |
| C | srlP | 3 | 40.76 |
| D | srlP | 1 | 41.61 |
| D | srlP | 2 | 41.06 |
| D | srlP | 3 | 40.85 |
| E | srlP | 1 | 41.16 |
| E | srlP | 2 | 41.72 |
| E | srlP | 3 | 40.81 |
| A | QIAseq | 1 | 32.04 |
| A | QIAseq | 2 | 30.60 |
| A | QIAseq | 3 | 31.90 |
| B | QIAseq | 1 | 32.18 |
| B | QIAseq | 2 | 32.22 |
| B | QIAseq | 3 | 32.70 |
| C | QIAseq | 1 | 32.49 |
| C | QIAseq | 2 | 31.94 |
| C | QIAseq | 3 | 31.41 |
| D | QIAseq | 1 | 31.25 |
| D | QIAseq | 2 | 32.14 |
| D | QIAseq | 3 | 31.19 |
| E | QIAseq | 1 | 32.62 |
| E | QIAseq | 2 | 33.00 |
| E | QIAseq | 3 | 32.19 |
| A | TailorMix | 1 | 34.92 |
| A | TailorMix | 2 | 35.14 |
| A | TailorMix | 3 | 35.99 |
| B | TailorMix | 1 | 34.87 |
| B | TailorMix | 2 | 35.07 |
| B | TailorMix | 3 | 35.39 |
| C | TailorMix | 1 | 34.94 |
| C | TailorMix | 2 | 35.67 |

| | | | |
|---|---|---|---|
| C | TailorMix | 3 | 34.93 |
| D | TailorMix | 1 | 35.08 |
| D | TailorMix | 2 | 35.26 |
| D | TailorMix | 3 | 35.47 |
| E | TailorMix | 1 | 36.46 |
| E | TailorMix | 2 | 37.47 |
| E | TailorMix | 3 | 36.89 |
| A | CleanTaq | 1 | 39.58 |
| A | CleanTaq | 2 | 39.53 |
| A | CleanTaq | 3 | 38.91 |
| B | CleanTaq | 1 | 39.61 |
| B | CleanTaq | 2 | 40.78 |
| B | CleanTaq | 3 | 41.22 |
| C | CleanTaq | 1 | 41.73 |
| C | CleanTaq | 2 | 42.09 |
| C | CleanTaq | 3 | 43.17 |
| D | CleanTaq | 1 | 41.97 |
| D | CleanTaq | 2 | 39.77 |
| D | CleanTaq | 3 | 42.27 |
| E | CleanTaq | 1 | 38.54 |
| E | CleanTaq | 2 | 40.88 |
| E | CleanTaq | 3 | 41.23 |

S8 Table

Table 8: Inter-rater correlations between rlog read counts and the theoretical sample concentration of the 40 non-equimolar miRNA oligonucleotides

| Kit | Mix | Replicate | Pearson correlation |
|---|---|---|---|
| CleanTag | A | 1 | 0.43 |
| CleanTag | A | 2 | 0.44 |
| CleanTag | A | 3 | 0.44 |
| CleanTag | B | 1 | 0.24 |
| CleanTag | B | 2 | 0.24 |
| CleanTag | B | 3 | 0.22 |
| CleanTag | C | 1 | 0.30 |
| CleanTag | C | 2 | 0.28 |
| CleanTag | C | 3 | 0.29 |
| CleanTag | D | 1 | 0.17 |
| CleanTag | D | 2 | 0.17 |
| CleanTag | D | 3 | 0.16 |
| CleanTag | E | 1 | 0.47 |
| CleanTag | E | 2 | 0.41 |
| CleanTag | E | 3 | 0.43 |
| QIAseq | A | 1 | 0.58 |
| QIAseq | A | 2 | 0.61 |
| QIAseq | A | 3 | 0.59 |
| QIAseq | B | 1 | 0.36 |
| QIAseq | B | 2 | 0.37 |
| QIAseq | B | 3 | 0.37 |
| QIAseq | C | 1 | 0.46 |
| QIAseq | C | 2 | 0.47 |
| QIAseq | C | 3 | 0.47 |
| QIAseq | D | 1 | 0.26 |
| QIAseq | D | 2 | 0.26 |
| QIAseq | D | 3 | 0.26 |
| QIAseq | E | 1 | 0.61 |
| QIAseq | E | 2 | 0.59 |
| QIAseq | E | 3 | 0.61 |
| srLp | A | 1 | 0.47 |
| srLp | A | 2 | 0.45 |
| srLp | A | 3 | 0.48 |
| srLp | B | 1 | 0.32 |
| srLp | B | 2 | 0.32 |
| srLp | B | 3 | 0.31 |
| srLp | C | 1 | 0.38 |
| srLp | C | 2 | 0.37 |

| | | | |
|---|---|---|---|
| srLp | C | 3 | 0.39 |
| srLp | D | 1 | 0.24 |
| srLp | D | 2 | 0.22 |
| srLp | D | 3 | 0.23 |
| srLp | E | 1 | 0.48 |
| srLp | E | 2 | 0.49 |
| srLp | E | 3 | 0.48 |
| TailorMix | A | 1 | 0.49 |
| TailorMix | A | 2 | 0.49 |
| TailorMix | A | 3 | 0.47 |
| TailorMix | B | 1 | 0.20 |
| TailorMix | B | 2 | 0.18 |
| TailorMix | B | 3 | 0.19 |
| TailorMix | C | 1 | 0.35 |
| TailorMix | C | 2 | 0.32 |
| TailorMix | C | 3 | 0.36 |
| TailorMix | D | 1 | 0.08 |
| TailorMix | D | 2 | 0.08 |
| TailorMix | D | 3 | 0.08 |
| TailorMix | E | 1 | 0.49 |
| TailorMix | E | 2 | 0.49 |
| TailorMix | E | 3 | 0.49 |
