## Supplementary Material and Methods for "Systematic assessment of commercially available low-input miRNA library preparation kits"

### Supplementary Materials & Methods

#### miRNA library preparation

##### Lexogen- srLp

In the commercially available miRNA library prep kit of Lexogen (srLp), Lexogen's Small RNA i7 index sequences are used. To minimise any possible bias arising due to the use of different index sequences, Lexogen agreed to use instead Illumina P7 index sequences as used by other participants in the study. Therefore, P7 PCR primers were exchanged for the PCR amplification step. All other steps were performed according to the user Guide available for download at https://www.lexogen.com/wp-content/uploads/2018/08/052UG128V0102_Small-RNA-Seq-Library-Prep-Kit.pdf. In detail, human total RNA samples were used with 1x A3/A5/RTP concentrations and 100 µl EtOH was added in step 5, while for the synthetic miRNA mixes A-E 0.5x A3/A5/RTP concentrations and 50 µl EtOH was added in step 5. Synthetic miRNA mixes A-E were amplified with 19 cycles, while total human RNA samples were amplified with 17 cycles. Synthetic miRNA mix libraries were only column purified according to the user guide post-PCR instructions. No further size selection was performed. For human total RNA samples, size selection (125 to 180 bp) was carried out using a 3 % agarose gel and Invitrogen's Purelink Quick Gel Extraction kit according to manufacturer's instructions. Elution volume was 50 µl; healthy control human total RNA samples were slightly below the desired concentration of 1ng/µl and were therefore concentrated by ethanol precipitation.

##### QIAGEN- QIAseq

NGS libraries were prepared from 10 ng and 1 ng synthetic miRNA and 100 ng human total RNA aliquots using the QIAseq miRNA Library Kit (QIAGEN) according to the manufacturer's handbook (https://www.qiagen.com/us/shop/sequencing/qiaseq-solutions/qiaseq-mirna-ngs/#resources). Briefly, adapters were ligated sequentially to the 3' (adapter diluted 1:5 for synthetic miRNA samples and undiluted for 100 ng human total RNA samples) and 5' (adapter diluted 1:2.5 for 10 ng synthetic miRNA samples, 1:2.5 for the 1 ng synthetic miRNA samples and undiluted for 100 ng human total RNA samples) ends of miRNAs. Subsequently, universal cDNA synthesis with unique molecular index (UMI) assignment (RT primer diluted 1:5 for 10 ng synthetic miRNA samples and undiluted for 100 ng human total RNA samples), cDNA cleanup, library amplification (16 cycles all samples) and library cleanup were performed per the manufacturer's handbook. In a first library preparation round, samples 1 to 8 of the synthetic miRNA revealed low library concentrations. A second set of samples was therefore shipped and library preparation repeated, which were subsequently returned for sequencing and data analysis in this paper.

##### SeqMatic- TailorMix

TailorMix miRNA Sample Preparation Kit (Version 3) was used to generate micro RNA libraries from the entire 5 µl of each RNA sample, according to the manufacturer's protocol (https://www.seqmatic.com/wp-content/uploads/2019/05/TM-310_-Workflow-U19D25001.pdf ). 15-PCR cycles were used during the PCR amplification step for all samples. Individual miRNA libraries were size-selected according to the manufacturer´s protocol (~140bp) and eluted in 30 µl TE buffer with 0.1% Tween-20. 10 µl of each eluted library was submitted for sequencing and data analysis in this paper.

#### TriLink-CleanTag

A detailed library preparation method can be found at https://www.trilinkbiotech.com/cleantag/ and Shore, et al. [1]. In brief, chemically modified 3' adapter was ligated to small RNA sample with T4 RNA Ligase 2 (KQ), followed by heat inactivation to stop the reaction. Chemically modified 5' adapter was then ligated to the library using T4 RNA Ligase 1. The reaction was heat inactivated and directly reverse transcribed without the need for purification using Protoscript II. The entire cDNA sample was then amplified with appropriate index primers by Q5 High Fidelity PCR master mix. For this work the following modifications to the protocol were applied: For the synthetic miRNA samples the CleanTag adapters were diluted 1:4 before use and 5µL of input sample per independent prep was used (the procedure accepts 2 – 10 µl RNA input volumes without adjustment to reagent volumes). For human total RNA samples, the CleanTag adapters were diluted 1:2 before use. At library amplification, 15 cycles of PCR was used for synthetic miRNA samples and 18 cycles of PCR for human total RNA samples. A 2-step AMPure bead based size selection purification was used after PCR with a 1x beads to sample volume followed by 1.8x beads to sample volume as detailed in the kit insert (https://www.trilinkbiotech.com/cart/coa/L3206_Insert.pdf). Sample 5 was returned at a concentration of 0.5 ng/µl instead of ≥1 ng/µl. However, the library was still suited for being sequenced and the results were used for the data analysis in this paper.

#### Takara- SMARTer

A user manual detailing the protocol for the released version of the kit is freely available for download at:

https://www.takarabio.com/assets/documents/User%20Manual/SMARTer%20microRNA-

Seq%20Kit%20User%20Manual_071018.pdf. To accommodate the increased input volume (5 ul vs the recommended 4 ul in the User Manual), some adjustments to the released protocol were applied as follows:

1) For the synthetic miRNA samples, 100ng of the 3' Adapter were used. For the human biological samples, the 3' Adapter was diluted to 1ng and then added to the 5 ul sample.

2) 12 ul, rather than 10 ul, of the Dephosphorylation reaction was transferred to the Circularization reaction and the volumes of the Circularization Master Mix were adjusted accordingly3) 12 ul, rather than 10 ul, of the Circularization reaction was used in the RT reaction and, again, the volumes of the RT reaction were adjusted accordingly.

Due to the lower than expected yield, the amplification cycles were increased (to 21 cycles) over what is listed in the kit protocol. Size selection was performed by Pippin Prep with a 3% gel, selecting the size range 128 bp to 160 bp.

##### Diagenode- CATS

All libraries were processed according to the protocol instructions as written in the CATS small RNA-seq kit (Diagenode, C05010040) manual (https://www.diagenode.com/files/products/kits/CATS-small-RNAseq-kit-manual.pdf). The standard protocol (step1: RNA de-phosphorylation and tailing) was employed for all small RNA library preparations, utilizing RT primer M for synthetic miRNA samples, and RT primer H for human total RNA samples. PCR amplification was performed for 9 cycles (sample mixes A-D), 12 cycles (mix E), or 10 cycles (human total RNA samples). After library amplification, the dsDNA solution was purified using the MicroChIP DiaPure spin columns as instructed in the manual (Diagenode, C03040001). The DNA was eluted in 20µl of EB buffer and was further processed in a gel-cut size selection of the miRNA-containing fragments. The size selection was performed as described in the supplementary protocol III: Precise Size Selection of DNA libraries with 4% E-Gel EX Agarose Gels of the CATS small RNA-seq kit except that the 4% agarose gel was crafted manually. After electrophoresis, the miRNA-containing band of the libraries was manually cut out and the DNA extracted as described in the protocol. Finally, the final size-selected libraries were quantified using the Qubit dsDNA HS quantification kit (ThermoFisher, Q32851) and the profiles assessed with the BioAnalyzer HS DNA assay (Agilent, 5067-4626). The CATS library preparation method was originally described in Turchinovich, et al. [2].

#### qPCR

Quantitative reverse-transcriptase PCR was performed using pre-designed TaqMan® Small RNA assays (Thermo Fisher Scientific, Waltham, MA USA) according to manufacturer's instructions. Assay details are provided in Table 1 below. cDNA was first prepared using the TaqMan MicroRNA Reverse Transcription Kit (Thermo Fisher Scientific), followed by qPCR with TaqMan Universal PCR Master Mix II (Thermo Fisher Scientific) on a QuantStudio 12K Flex instrument (Thermo Fisher Scientific). Relative abundances of miRNAs in mixes A and B were measured by absolute quantification relative to a standard curve.

Table 1: TaqMan Small RNA assay details

| miRNA | Assay ID | miRNA sequence | Cat # |
|---|---|---|---|
| hsa-miR-380-5p | 570 | UGGUUGACCAUAGAACAUGCGC | 4427975 |
| hsa-miR1224-3p | 2752 | CCCCACCUCCUCUCUCCUCAG | 4427975 |
| hsa-miR-136-3p | 2100 | CAUCAUCGUCUCAAAUGAGUCU | 4427975 |
| hsa-miR-30c-2-3p | 2110 | CUGGGAGAAGGCUGUUUACUCU | 4427975 |
| hsa-miR-652-3p | 2352 | AAUGGCGCCACUAGGGUUGUG | 4427975 |
| hsa-miR-15b-3p | 2173 | CGAAUCAUUAUUUGCUGCUCUA | 4427975 |

| | | | |
|---|---|---|---|
| hsa-let-7i-3p | 2172 | GGGGCUGGGGCCGGGACAGAGC | 4427975 |
| hsa-miR-374a-3p | 2125 | CUUAUCAGAUUGUAUUGUAAUU | 4427975 |
| hsa-miR-22-5p | 2301 | AGUUCUUCAGUGGCAAGCUUUA | 4427975 |
| hsa-miR-384 | 574 | AUUCCUAGAAAUUGUUCAUA | 4427975 |
| *hsa-miR-99b-3p* | 2196 | CAAGCUCGUGUCUGUGGGUCCG | 4427975 |
| hsa-miR-496 | *1953* | UGAGUAUUACAUGGCCAAUCUC | 4427975 |
| *hsa-miR-545-5p* | 2266 | UCAGUAAAUGUUUAUUAGAUGA | 4427975 |
| *hsa-miR-1915-3p* | 121111_mat | CCCCAGGGCGACGCGGCGGG | 4427975 |
| hsa-miR-125b-1-3p | 2378 | ACGGGUUAGGCUCUUGGGAGCU | 4427975 |
| hsa-miR-711 | 241090_mat | GGGACCCAGGGAGAGACGUAAG | 4427975 |

#### Trimming

Cutadapt version 1.15 was used for trimming 3' adapter sequences with the option –m 10. The following adapter sequences or commands were used: (1) adapter sequence for TailorMix and CleanTag: TGGAATTCTCGGGTGCCAAGG, (2) adapter sequence for SMARTer: TGGAATTCTCGGGTGCCAAGGC

(3) adapter sequence for srLp: TGGAATTCTCGGGTGCCAAGGAACTCCAGTCAC, (4) Adapter sequence for QIAseq: AACTGTAGGCACCATCAAT, (5) adapter trimming command for CATS: AATTCTCGGGTGCCAAGGAACTC

cutadapt -u 3 input_file.fastq | cutadapt -a AAAAAAAA - | cutadapt -a AAAAAAAN$ -a AAAAAAN$ -a AAAAAN$ - | cutadapt -a AGAGCACACGTCTG - | cutadapt -O 8 -g GTTCAGAGTTCTACAGTCCGACGATCNNN | cutadapt -m 10 -o output.file.
